# Sulfonylureas target the neurovascular response to decrease Alzheimer’s pathology

**DOI:** 10.1101/2021.08.11.455969

**Authors:** Shannon L. Macauley, Molly S. Stanley, Emily E. Caesar, William R. Moritz, Annie R. Bice, Nildris Cruz-Diaz, Caitlin M. Carroll, Stephen M. Day, John Grizzanti, Thomas E. Mahan, James A. Snipes, Timothy E. Orr, Joseph P. Culver, Maria S. Remedi, Colin G. Nichols, Celeste M. Karch, Laura A. Cox, Debra I. Diz, Adam Q. Bauer, David M. Holtzman

## Abstract

Hyperexcitability is a defining feature of Alzheimer’s disease (AD), where aberrant neuronal activity is both a cause and consequence of AD. Therefore, identifying novel targets that modulate cellular excitability is an important strategy for treating AD. ATP-sensitive potassium (K_ATP_) channels are metabolic sensors that modulate cellular excitability. Sulfonylureas are K_ATP_ channel antagonists traditionally used to combat hyperglycemia in diabetic patients by inhibiting pancreatic K_ATP_ channels, thereby stimulating insulin release. However, K_ATP_ channels are not limited to the pancreas and systemic modulation of K_ATP_ channels has pleotropic physiological effects, including profound effects on vascular function. Here, we demonstrate that human AD patients have higher cortical expression of vascular K_ATP_ channels, important modulators of vasoreactivity. We demonstrate that peripheral treatment with the sulfonylurea and K_ATP_ channel inhibitor, glyburide, reduced the aggregation and activity-dependent production of amyloid-beta (Aβ), a hallmark of AD, in mice. Since glyburide does not readily cross the blood brain barrier, our data suggests that glyburide targets vascular K_ATP_ channel activity to reduce arterial stiffness, improve vasoreactivity, and normalize pericyte-endothelial cell morphology, offering a novel therapeutic target for AD.

**Graphical abstract:** Targeting vascular K_ATP_ channel activity for the treatment of Alzheimer’s disease pathology.

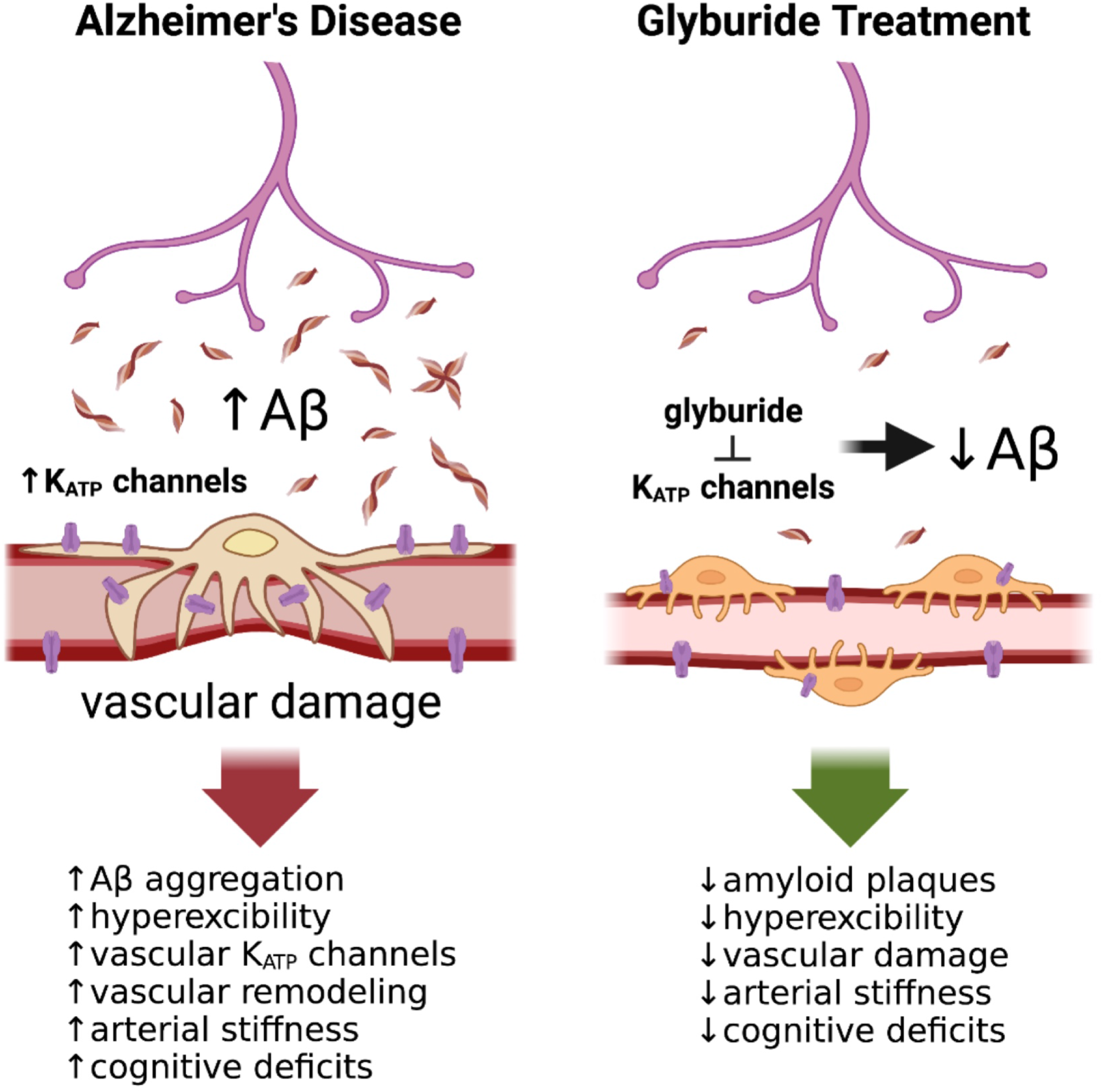

## INTRODUCTION

Normal cellular excitability underlies physiological function and when altered, can lead to or exacerbate disease. Brain hyperexcitability is central to the pathophysiology of Alzheimer’s disease, with aberrant neuronal activity is both a cause and consequence of Alzheimer’s pathology. Neuronal release of amyloid-β (Aβ) and tau, the pathological hallmarks of AD, is driven by changes in neuronal activity [1–7]. Neurons generate soluble, monomeric Aβ via the sequential cleavage of the amyloid precursor protein (APP) and secrete Aβ into the extracellular space [8, 9]. Aβ release at the synapse occurs in an activity dependent manner, whereby increased activity raises interstitial fluid (ISF) Aβ levels, which can then form toxic, higher order species and aggregate into amyloid plaques in a concentration-dependent manner [1, 2, 10, 11]. Thus, increasing neuronal excitability will increase extracellular Aβ release and promote plaque formation in the brain. Similarly, targeting aberrant excitability could mitigate Aβ release and amyloid plaque development.

Widely distributed in excitable cells, ATP-sensitive potassium (K_ATP_) channels link changes in metabolism with electrical activity [12, 13]. In pancreatic beta cells, K_ATP_ channels close in response to rising blood glucose levels, triggering insulin secretion. K_ATP_ channels are also expressed in multiple neuronal subtypes. Our previous studies demonstrate that pharmacological manipulation of neuronal K_ATP_ channels altered neuronal excitability and ISF Aβ levels [7]. This work echoed previous findings that increased cerebral metabolism promotes neuronal K_ATP_ channel closure and increased excitability [14–16]. In vascular smooth muscle cells, K_ATP_ channels regulate blood flow, vasodilation/vasoconstriction, and vascular tone [12, 17, 18]. Since hyperexcitability is a key feature of AD, we hypothesized that changes in K_ATP_ channel activity at the neurovascular unit can alter neuronal activity and AD pathogenesis, and hence, serve as a therapeutic target in AD.

Epidemiological studies suggest that sulfonylureas, a class of FDA approved drugs that inhibit K_ATP_ channel activity [19], may reduce AD and dementia risk, but the mechanisms are unknown [20–22]. To date, no prospective study in humans has evaluated the effects of sulfonylureas on vascular health, brain function, or cognition, in AD. Interestingly, sulfonylureas are widely used to treat type 2 diabetes, a risk factor for AD [23], and currently being assessed in preclinical and clinical studies for other CNS diseases, including stroke and traumatic brain injury [24–27].

The goal of this work was to elucidate the role of vascular K_ATP_ channels in AD and determine whether K_ATP_ channel inhibition with glyburide could alter AD pathogenesis. We find that K_ATP_ channel subunits localized to vascular smooth muscle cells, endothelial cells, and/or pericytes are upregulated in human AD brains. Using an established model of cerebral amyloidosis, the APPswe/PSEN1dE9 (APP/PS1) mouse [28], we demonstrate that peripheral administration of the sulfonylurea glyburide decreased amyloid burden and altered stimulus-driven neural and hemodynamic activity. Consistent with Aβ being released in an activity dependent manner, we further demonstrate that decreased evoked responses lead to decreased hippocampal ISF Aβ levels. Ultimately, these reductions were associated with decreased Aβ aggregation. Given that glyburide does not efficiently cross the blood-brain barrier (BBB) [29] and that vascular K_ATP_ channels regulate vasoreactivity, we explored whether glyburide treatment impacted cerebrovascular morphology and function. In mice, glyburide altered the neurovascular response, reduced arterial stiffness, and normalized pericyte-endothelial cell morphology. Short term glyburide treatment also improved learning and memory in APP/PS1 mice. Taken together, these data suggest that K_ATP_ channels not only play a critical and previously unappreciated role linking Aβ with neurovascular dysfunction in AD, but also offer a novel therapeutic target in AD.

## RESULTS

### K_ATP_ channels are upregulated in human Alzheimer’s disease brains

Using the publicly available database from Mayo RNAseq Study (via AMP-AD Knowledge Portal), we investigated K_ATP_ channel expression in the temporal cortex in pathological aging (PA=amyloid without tau), Alzheimer’s disease (AD=amyloid and tau), and progressive supranuclear palsy (PSP=tau without amyloid) (Fig 1). *KCNJ8, KCNJ11, ABCC8, ABCC9* genes code for the K_ATP_ channel subunits, Kir6.1, Kir6.2, SUR1, and SUR2, respectively. At autopsy, Kir6.1 (*KCNJ8*) and SUR2 (*ABCC9*) are both significantly upregulated in AD, but not in NC, PA, or PSP. Kir6.1 and SUR2 heterodimerize to form K_ATP_ channels localized to mural cells, vascular smooth muscle cells, endothelial cells, and pericytes [30], and *ABCC9* is widely used as a transcriptional signature for pericytes [31–33]. No significant changes were observed in *KCNJ11* or *ABCC8* in any conditions (q value < 0.05). These subunits are typically enriched in neurons [34], astrocytes [35], and oligodendrocytes [16]. These results thus indicate that vascular, but not neuronal, K_ATP_ channel expression is specifically increased in human AD cortex, but not in other neurodegenerative diseases.

**Figure 1.**
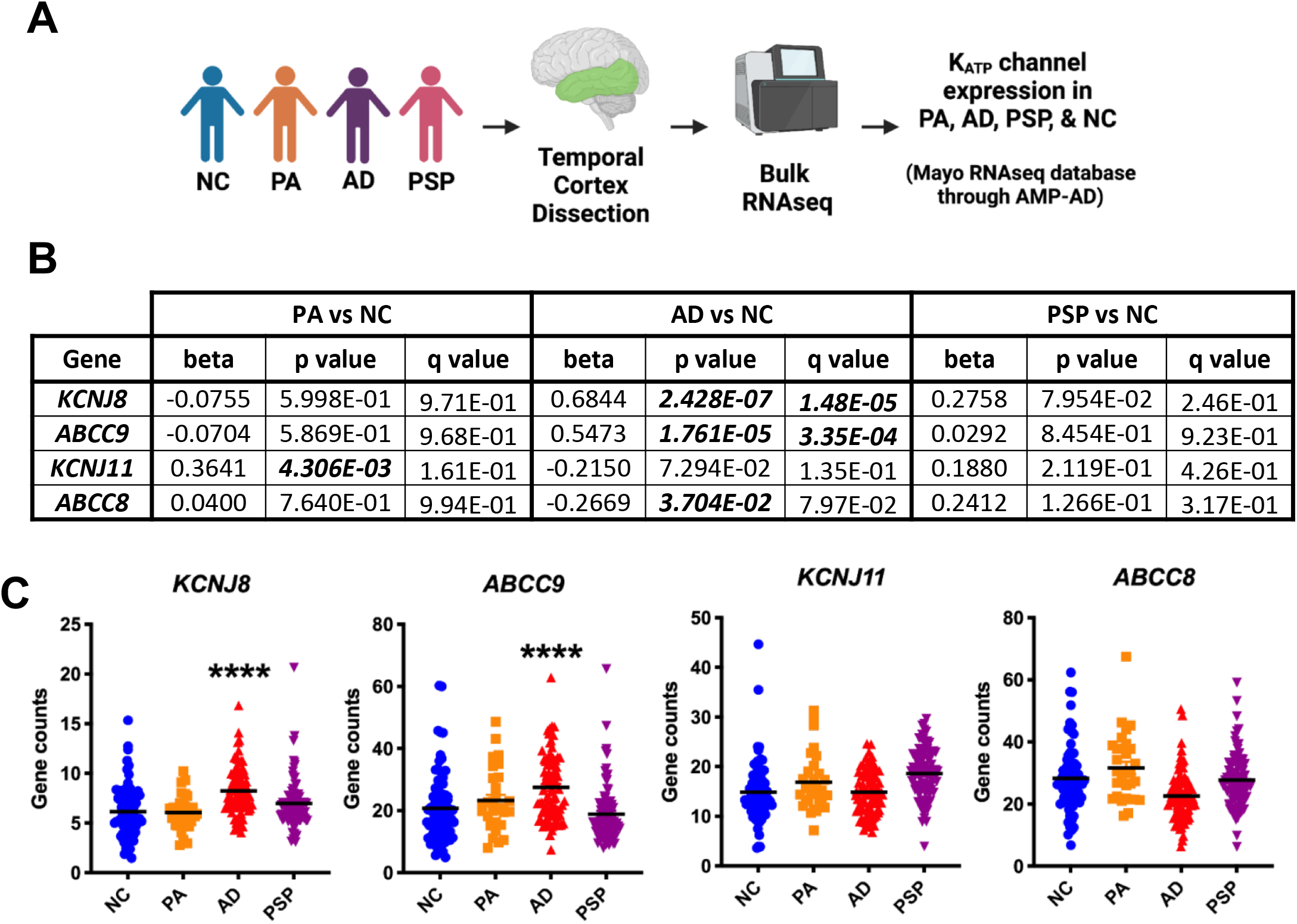
Increased vascular K_ATP_ channel expression in the temporal cortex of Alzheimer’s disease patients at autopsy. (A) Workflow to explore how K_ATP_ channel genes (*ABCC8, ABCC9, KCNJ8, KCNJ11*) in the temporal cortex change due to AD-related pathology using the Mayo RNAseq database. (B&C) Increased K_ATP_ channel subunit expression in the cortex of Alzheimer’s patients (AD), but not in normal controls (NC), pathological aging (PA), or progressive supranuclear palsy (PSP). Significant changes (q value < 0.05; bolded and italicized) were observed in *KCNJ8* (q value=2.43E-07) and *ABCC9* (q value=1.76E-05), but not in *KCNJ11* and *ABCC8. KCNJ8* and *ABCC9* heterodimerize to form K_ATP_ channels on the vasculature, including mural cells, pericytes, and endothelial cells.

### Systemic glyburide treatment reduces Aβ deposition and plaque pathology in APP/PS1 mice

Given the increased K_ATP_ channel expression in AD brains, we explored whether targeting these channels therapeutically might impact AD-related pathology. Four-month-old female APP/PS1 mice were treated with glyburide or placebo for 3 months via subcutaneous implanted slow release pellet (2.5mg/pellet; 30μg/day)[29, 36]. Representative images from placebo-treated mice show prominent plaque pathology in the cortex and hippocampus, but reduced Aβ deposition in glyburide-treated mice (Fig 2A). When quantified, Aβ deposition decreased by ~50% (Fig 2B) and amyloid plaques decreased by ~45% (Fig 2C) in both cortex and hippocampus. Insoluble levels of Aβ40 and Aβ42 decreased by ~30-40% in the hippocampus of glyburide-treated animals (Fig 2D), while soluble levels for both Aβ40 and Aβ42 remained unchanged. Taken together, these experiments demonstrate that systemic glyburide treatment reduces Aβ pathology and amyloid plaque burden.

**Figure 2.**
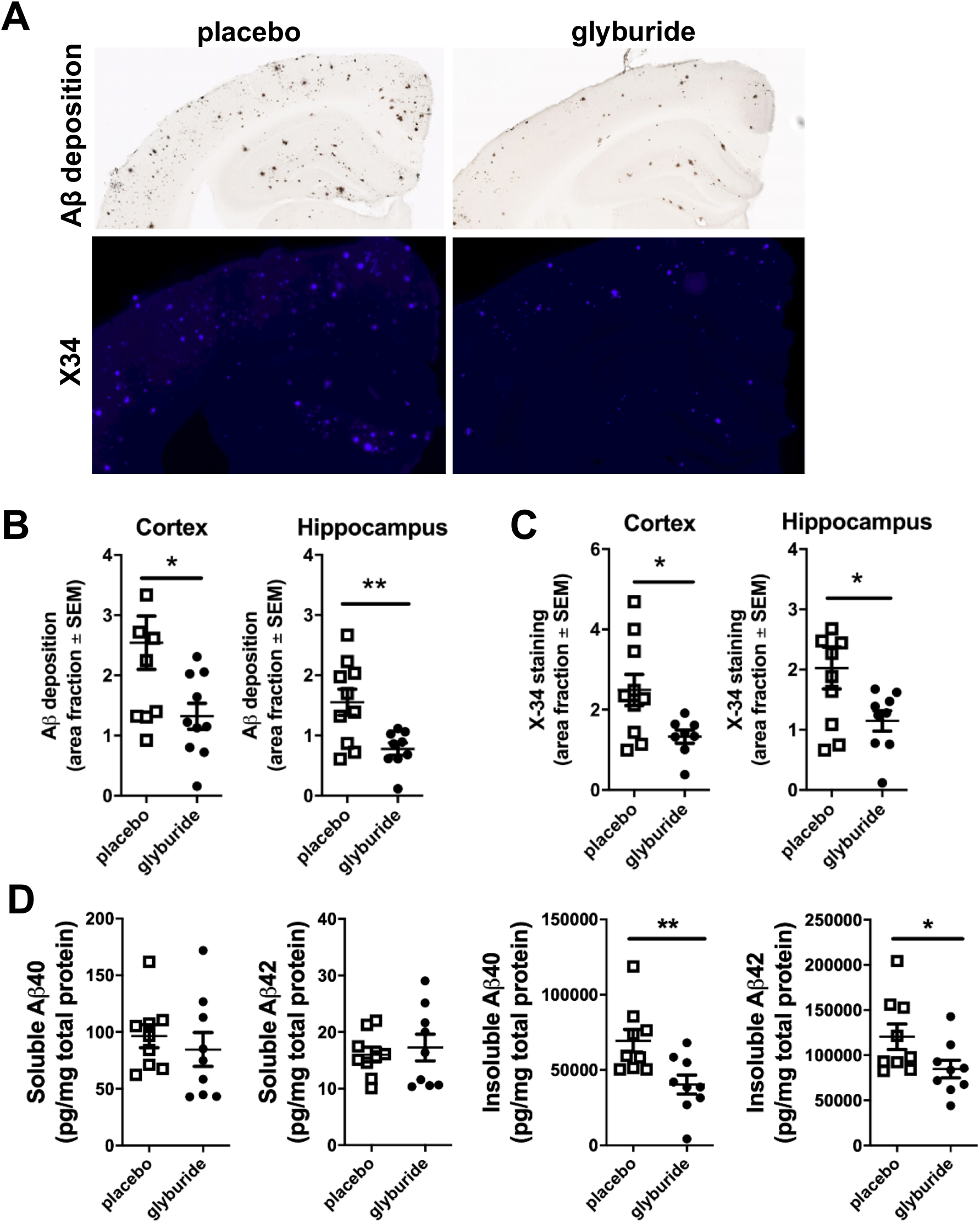
Systemic treatment with the sulfonylurea, glyburide, reduces Aβ and plaque pathology in APP/PS1 mice. (A) Representative images from the cortex and hippocampus of glyburide and placebo treated mice stained for Aβ deposition (HJ3.4B) and plaque pathology (X34; n=9-10 mice/group). (B) Aβ deposition decreased in glyburide treated mice compared to placebo in the cortex and hippocampus by 48% and 50%, respectively. (C) Plaque pathology decreased in glyburide treated mice compared to placebo in both the cortex and hippocampus by 47% and 43%, respectively. (D) The insoluble levels of Aβ40 and Aβ42 decreased by 42% and 30%, respectively, in hippocampus of glyburide treatment mice. Significance determined via t-test with *p<0.05; **p<0.001.

### Systemic glyburide treatment does not affect Aβ clearance mechanisms

While glyburide treatment reduced Aβ deposition, amyloid plaque pathology, and insoluble Aβ40 and Aβ42, it is unclear whether this was the result of decreased Aβ production or increased Aβ clearance. Therefore, we explored Aβ clearance mechanisms (Fig 3&4). First, we administered compound E, a potent gamma secretase inhibitor, to glyburide- or placebo-treated mice to prevent amyloid precursor protein (APP) cleavage and Aβ production, then quantified Aβ clearance from the hippocampal ISF. No difference in Aβ_1-x_ half-life was observed between glyburide- and placebo-treated mice (Fig 3A). There were also no differences in mRNA levels for Aβ degrading enzymes, insulin degrading enzyme (*Ide*) or neprilysin (*Nep*) (Fig 3B), nor in IDE protein levels. Similarly, no difference in the expression levels of clearance related proteins, such as *Apoe, Lrp1, Aqp4 or Rage,* was detected between glyburide or placebo groups (Fig 3C).

**Figure 3.**
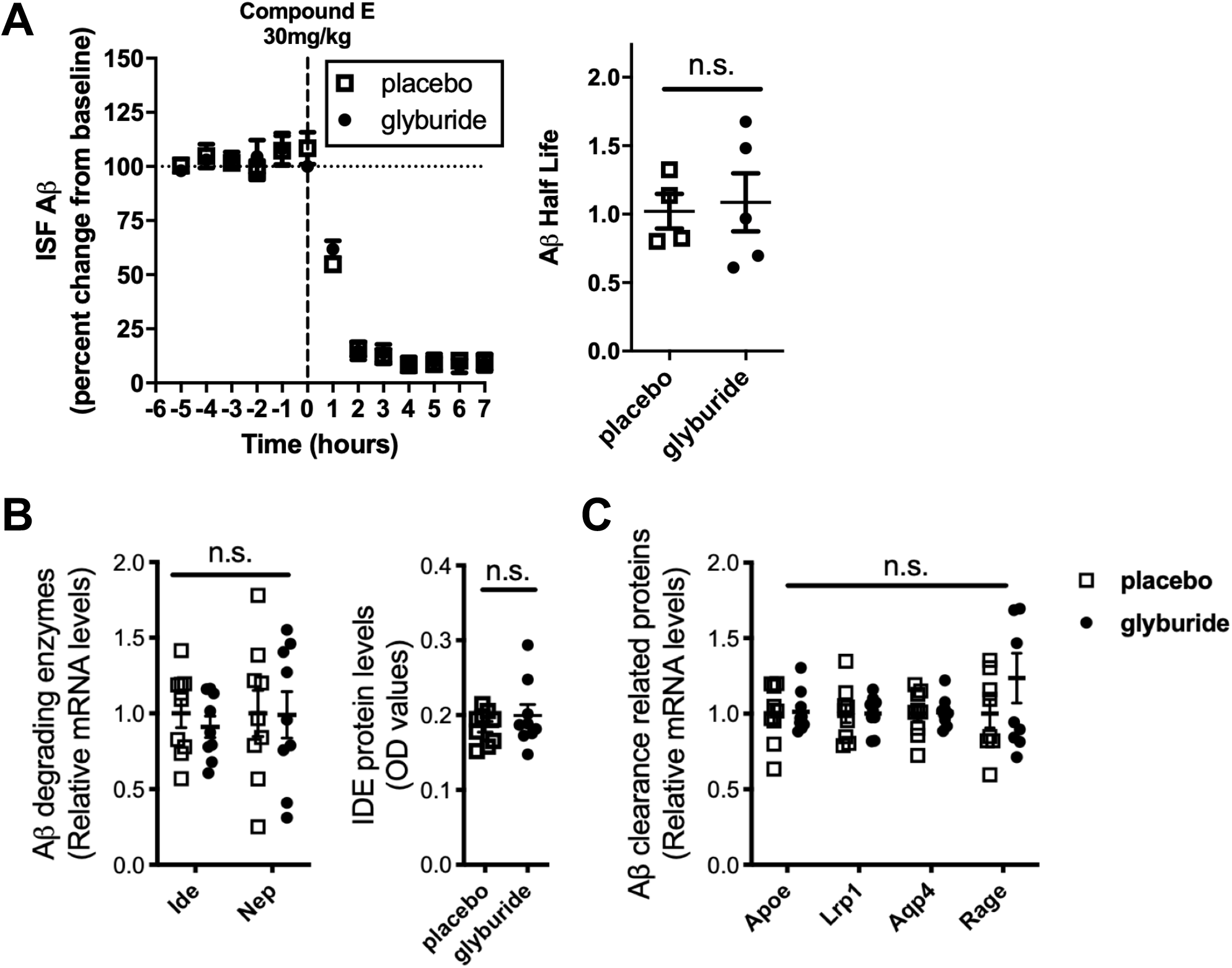
Systemic glyburide treatment does not affect Aβ clearance mechanisms. (A) There was no difference in Aβ half-life following treatment with a gamma secretase inhibitor, compound E, in glyburide treated mice compared to placebo suggesting Aß clearance is not affect by glyburide treatment (n=4-5 mice/group). (B) There was no difference in Aβ degrading enzymes, insulin degrading enzyme (*Ide*) or neprilysin (*Nep*), at either the transcript or protein level (n=9 mice/group). (C) No change in the mRNA levels of clearance related proteins, such as *Apoe, Lrp1, Aqp4, or Rage* (n=9 mice/group). Significance determined via t-test with n.s.= not significant.

**Figure 4.**
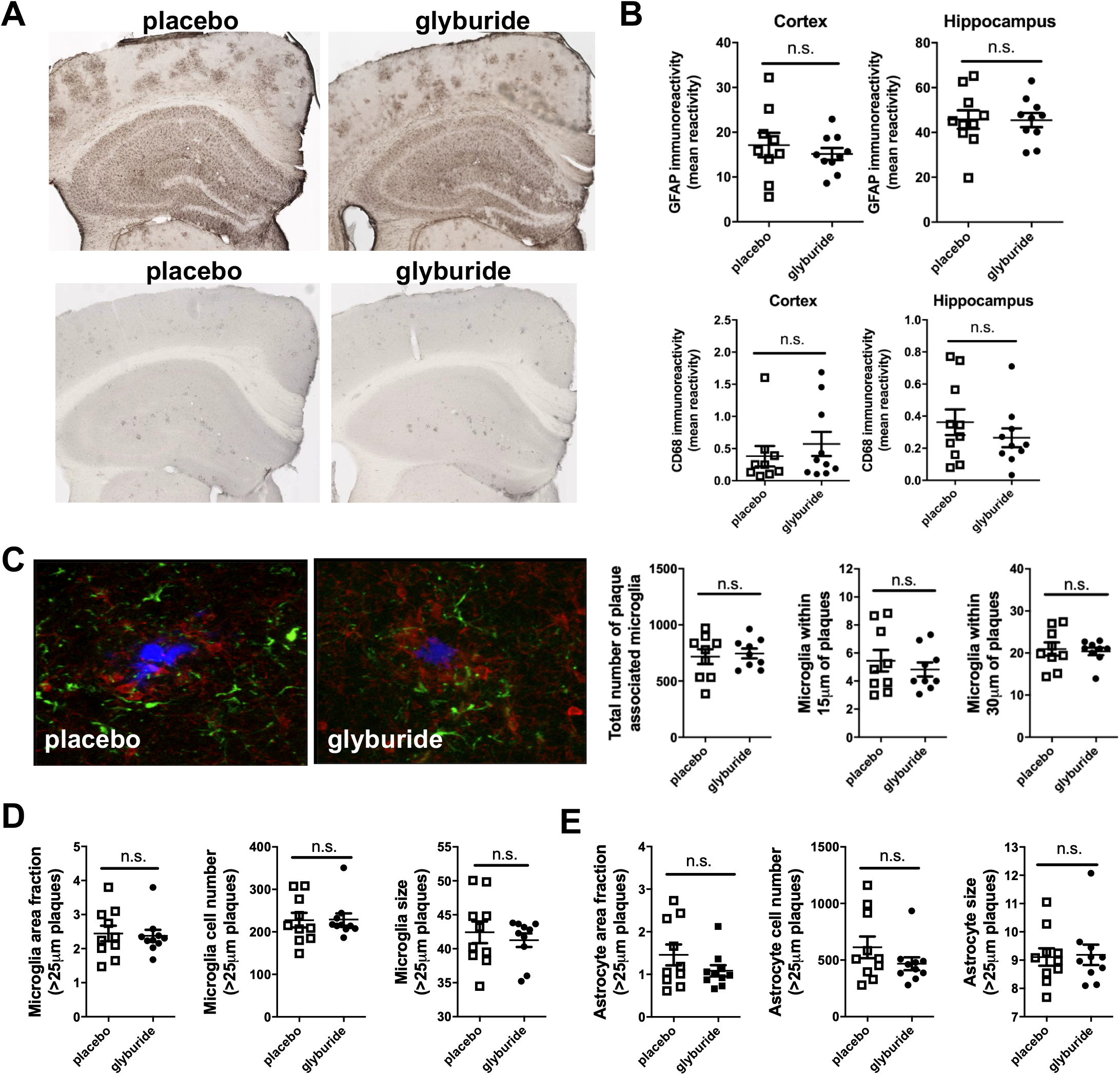
Neuroinflammation is unchanged in glyburide treated mice compared to controls despite reduced Aβ deposition. (A) Astroctye activation (GFAP staining) was comparable in the cortex and hippocampus of glyburide- and placebo-treated mice (n=9-10 mice/group). (B) Levels of CD68 were comparable in glyburide group compared to placebo. (C) Triple labeling of mouse cortex for amyloid plaques (X-34, blue), astrocytes (GFAP, green), and microglia (Iba1, red) in glyburide and placebo treated mice. (D) No difference in area fraction, total number, or average size of microglia or astrocytes. (E) No difference in number of plaque associated microglia between placebo and glyburide treated mice (amyloid plaques=X34; blue, astrocytes=GFAP; green, and microglia=CD68; red). Significance determined via t-test with n.s.= not significant.

Upregulation of the neuroinflammatory milieu, in particular astrocyte and microglial activation, also serves as a potential clearance mechanism for Aβ, and typically co-varies with amyloid plaque deposition. Therefore, glyburide- and placebo-treated brains were immunostained for GFAP and CD68, markers of astrocyte and microglial reactivity, respectively (Fig 4). Comparable levels of activated astrocytes (Fig 4A) and microglia (Fig 4B) were observed in the cortex and hippocampus of treated mice compared to controls at 7-months of age. Triple labeling of the mouse cortex for amyloid plaques (X34; blue), astrocytes (GFAP; green), and microglia (CD68; red) was performed to assess glial activation either proximal or distal to plaque pathology (Fig 4C). There was no difference in the total number of plaque-associated microglia nor in the number of microglia 15um or 30 um from the plaques (Fig 4C). Similarly, at distances greater than 25um from the plaques, there was no difference in the area fraction, total number, or size of activated microglia or astrocytes, suggesting comparable levels of glial activation in the glyburide- and placebo-treated mice (Fig 4D-E). Taken together, these data indicate that Aβ clearance pathways are unaltered in glyburide-treated mice.

### Glyburide reduces activity dependent release of Aβ by altering the neurovascular response

Aβ is produced via synaptic activity, where increased neuronal activity drives Aβ release into the ISF [10, 37]. Therefore, we assessed activity dependent release of Aβ in placebo- and glyburide-treated animals, by monitoring neuronal activity, ISF Aβ levels, and amyloid precursor protein (APP) processing. First, EEG recordings were performed in forelimb somatosensory cortex (S1FL) during electrical stimulation of the contralateral forepaw (Fig 5). During the evoked response, the placebo group exhibited a robust (~40%) increase in EEG amplitude, while glyburide treated mice did not (~10%) (Fig 5A, B, D). Interestingly, resting state EEGs and steady state ISF lactate levels, a marker of neuronal activity, were similar between glyburide and placebo groups, suggesting that basal neuronal activity was unaffected (Fig 5C, F). Next, we investigated ISF Aβ levels during the resting state and after administration of picrotoxin (PTX), a GABA antagonist, to stimulate excitatory neurotransmission [1, 2, 10, 38]. Steady state levels of ISF Aβ were ~25% lower in the glyburide-treated group versus placebo (Fig 5E). ISF lactate, a marker of neuronal activity, increased in both the glyburide- and placebo-groups after PTX administration, confirming enhanced neuronal activity. Interestingly, PTX increased ISF Aβ in the placebo-, but not the glyburide-treated group (Fig 5F). This indicates that ISF Aβ levels were reduced by glyburide treatment during the resting state and the evoked response in APP/PS1 mice. Since PTX also induces seizures, we monitored seizure activity during the 2 hrs post-PTX administration. Although seizure number was similar between groups, there was a significant reduction in the variance in the glyburide-treated group, suggesting glyburide influenced hyperexcitability in the APP/PS1 mice. Lastly, we explored whether APP processing and Aβ metabolism differed between groups. APP levels were comparable between glyburide- and placebo-treated mice (Fig 5H), although there was a reduction in amyloidogenic processing of APP in the glyburide-treated. Taken together, the results indicate that systemic glyburide treatment reduces the activity dependent release of Aβ while preserving resting state neuronal activity.

**Figure 5.**
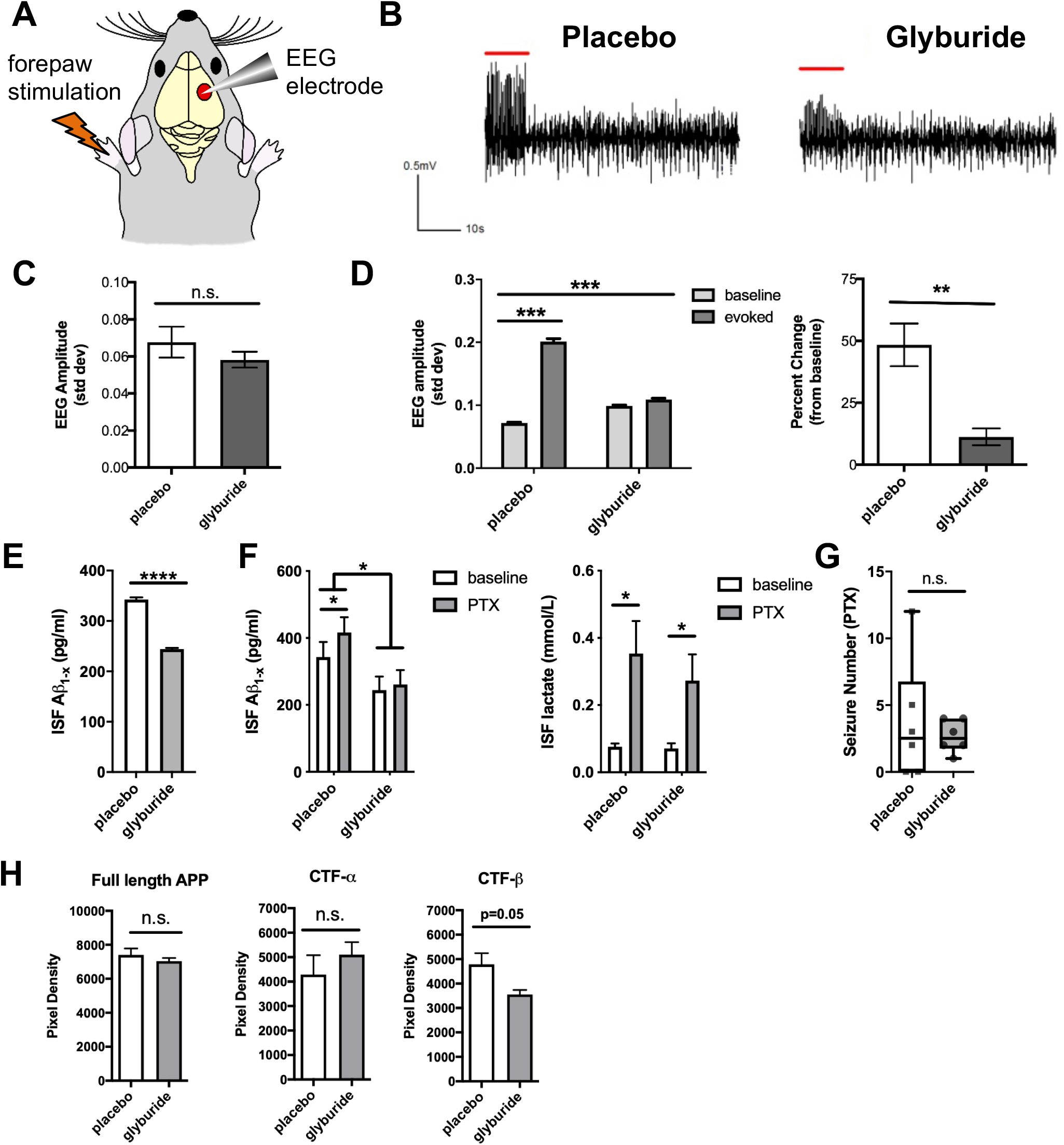
Reduced evoked responses and ISF Aβ levels in glyburide treatment mice. (A) Left forepaw stimulation was paired with EEG recordings in the contralateral somatosensory cortex (S1BL). (B) Representative EEG traces from S1FL cortex during 10 seconds of stimulation followed by 50 seconds of rest. (C) Basal EEGs are similar between placebo and glyburide groups (n=5 mice/group). (D) During forepaw stimulation, EEG amplitude increased by ~40% in placebo group, but only 10% in glyburide group. (E) Steady state levels of ISF Aβ are lower in glyburide group compared to placebo. (F) PTX treatment increased ISF Aβ levels in placebo but not in glyburide group, while ISF lactate, a marker of neuronal activity, increased in both (n=6/group). (G) No difference in seizure frequency between groups post-PTX but variance in glyburide group decreased (n=6 mice/group). (H) Decreased amyloidogenic processing of APP was seen in glyburide mice (n=4-5 mice/group). Significance determined via t-test, one-way ANOVA, or two-way ANOVA with Tukey’s post hoc tests when appropriate. *p<0.05, **p<0.01, ***p<0.001, ****p<0.0001

Since glyburide does not efficiently cross the BBB [29], a direct action on neurons and glia is unlikely. Instead, we hypothesize that glyburide affects vascular K_ATP_ channel activity to alter the neurovascular response. We therefore investigated whether glyburide affected evoked hemodynamic responses in APP/PS1 mice (Fig 6). Following forepaw stimulation, both placebo- and glyburide-treated groups exhibited stimulus evoked responses consistent with functional hyperemia: increased CBF, HbO, HbT, CMRO2 and decreased HbR (Fig 6A). However, consistent with observations from electrophysiology experiments (Fig 5B), evoked changes in CBF, HbO, and HbT due to stimulation were all significantly lower in the glyburide-treated group compared to placebo (Fig 6B). Although not significant, OEF was proportionately increased, and steady state levels of ISF glucose and ISF lactate, the two main energy substrates for the brain, were comparable between glyburide and placebo treated mice (Fig 6C). Nor was there any perturbation of peripheral metabolism with glyburide treatment, further suggesting that alterations in hemodynamic activity did not negatively affect cerebral or peripheral metabolism (Fig 6D).

**Figure 6.**
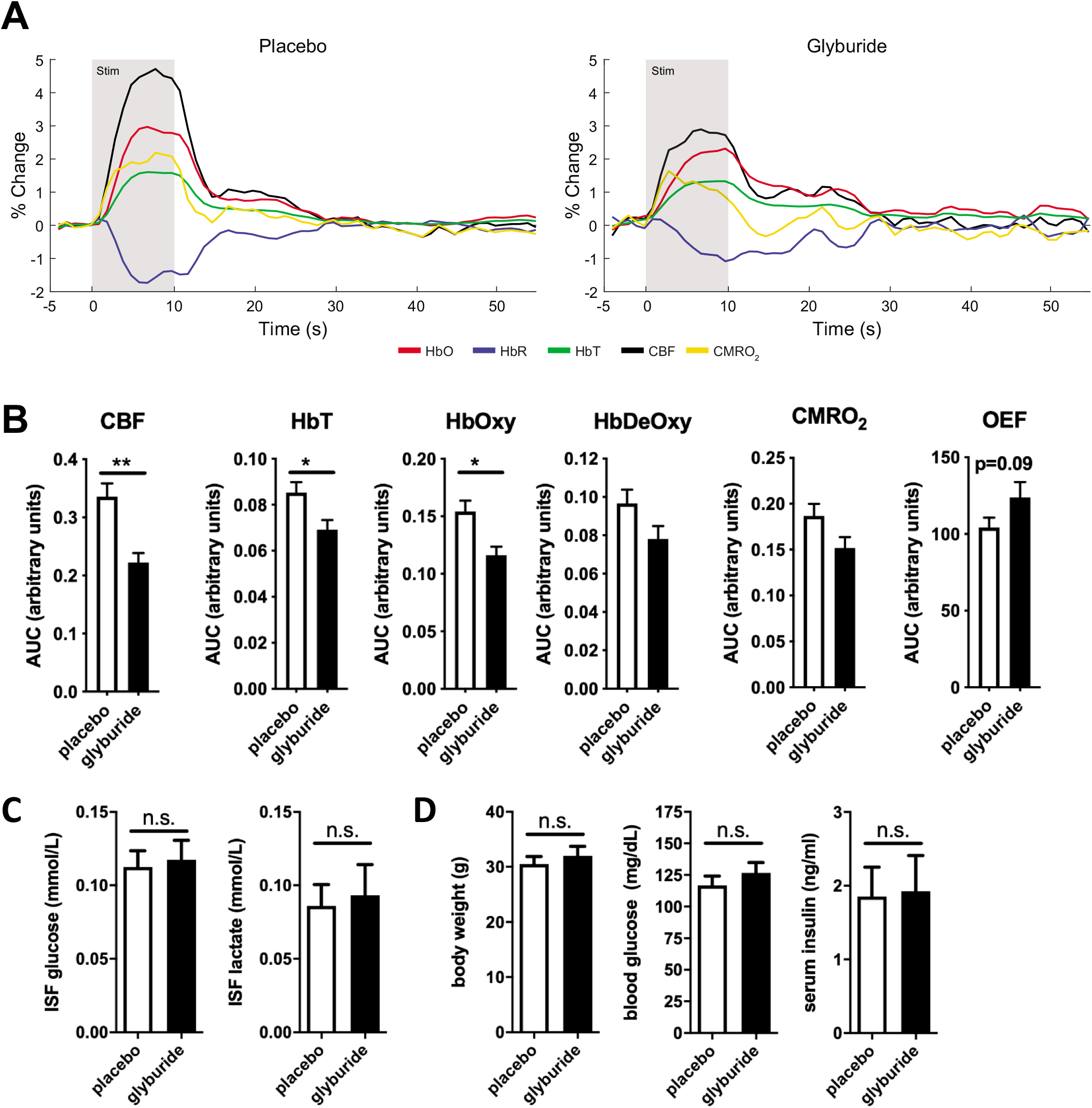
Glyburide treatment decreased stimulus-evoked hemodynamic activity in the cortex without altering cerebral or peripheral metabolism in APP/PS1 mice. (A) Group-averaged evoked time courses of oxygenated (HbOxy), Deoxygenated (HbDeOxy) and total (HbT) hemoglobin, cerebral blood flow (CBF), cerebral metabolic rate of oxygen consumption (CMRO2), and oxygen extraction fraction (OEF) following forepaw stimulation (n=8-9 mice/group). (B) Significant differences were observed in the responses for CBF, HbOxy, and HbT, but CMRO2 was preserved due to increased OEF in glyburide mice. (C) Steady state levels of ISF glucose and lactate were similar in placebo and glyburide groups (n=4-6 mice/group). (D) Body weight, blood glucose levels, and serum insulin levels were comparable in placebo and glyburide groups (n=9-10 mice/group). Significance determined via t-test with *p<0.05; **p<0.001.

Given the differences in hemodynamic activity observed between groups, we explored the possibility that glyburide treatment might cause vascular remodeling in the APP/PS1 brain. Since K_ATP_ channels are found throughout the vasculature, including endothelial cells and pericytes, we explored whether glyburide affected pericyte-endothelial cell morphology (Fig 7). Staining with CD31 (endothelial cells) and CD13 (pericytes) demonstrated that CD31 and CD13 were upregulated in the cortex of 7-month-old APP/PS1 compared to WT controls. Interestingly, glyburide-treatment of APP/PS1 mice reduced CD31 and CD13 staining, similar to WT levels (Fig 7C). Vessel diameter and width were increased in APP/PS1 mice, but this effect was reversed in glyburide treated mice (Fig 7C). This suggests that chronic treatment with glyburide prevents vascular remodeling caused by Aβ pathology.

**Figure 7.**
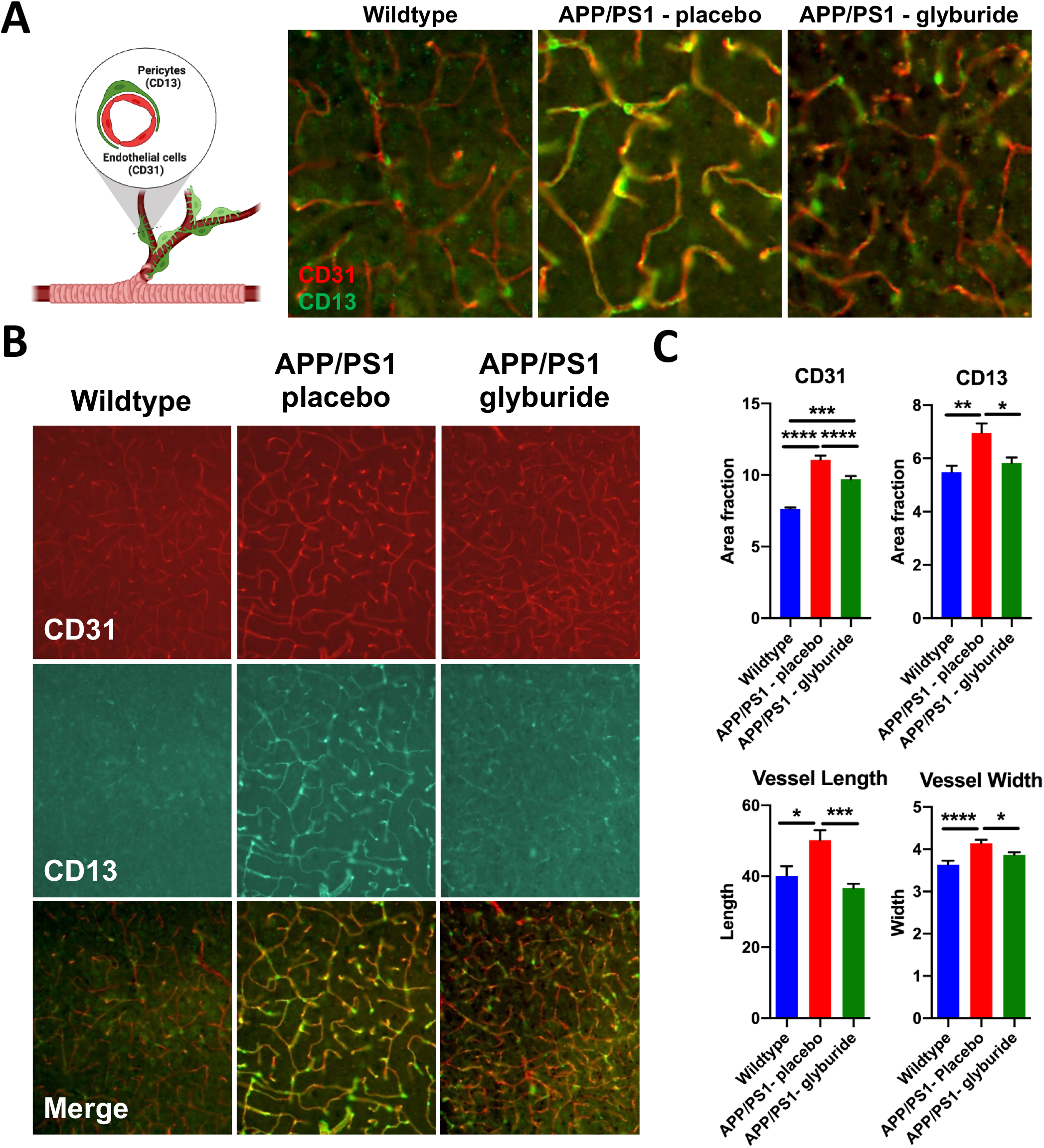
Glyburide restores pericyte-endothelial cell morphology. (A) High magnification image of CD31 (red; endothelial cells) and CD13 (green; pericytes) in wildtype, APP/PS1 placebo, and APP/PS1 glyburide groups (n=9-10/group). (B) Low magnification of CD31, CD13, and merged images from wildtype, APP/PS1 placebo, and APP/PS1 glyburide groups. (C) Quantification of CD31, CD13, vessel length, and vessel width in all groups. APP/PS1 placebo group has increased CD31 and CD13 staining as well as vessel width and vessel length in the cortex compared to wildtype mice. Glyburide treatment reduced all metrics to wildtype levels. Significance determined via one-way ANOVA with Tukey’s posthoc tests with *p<0.05; **p<0.001.*p<0.05, **p<0.01, ***p<0.001, ****p<0.0001.

### Glyburide treatment reverses canonical pathways associated with vessel hypertrophy and cellular excitability

Microarray analysis of gene expression was performed on cortical tissue extracted from APP/PS1 and WT mice following glyburide or placebo treatment to assess disease- or treatment-specific changes (Figure 8). First, differentially expressed genes (DEGs) were identified using Bayesian statistics; significant genes were filtered out by a p value<0.05, FDR-q<0.05, and a fold-change>/=2. We found 1576 genes were differentially expressed in APP/PS1 brains compared to WT, with 851 genes upregulated and 725 genes downregulated. In the glyburide-treated APP/PS1 group compared to placebo treated APP/PS1 group, 341 genes were differentially expressed, with 161 genes being upregulated and 180 genes being downregulated (Figure 8A). 32 genes were differentially expressed in APP/PS1 v WT mice and reverse in glyburide-treated v placebo-treated APP/PS1 mice (Figure 8A). Of those 32 DEGs, 23 DEGs had known gene names (Figure 8B). The 23 DEGs were enriched for pathways including regulation of G protein-coupled receptor activity and signal transduction (e.g *Ffar4, Ect2l, Tas2r138, Vmn1r234, Lpcat2*), protein modification and proteolysis (e.g. *Adam18, Metap1d, Pcsk5, Pld5, Zar1*), gene expression (e.g. *Dnase1l1, Ddmt3a, Nova1*), redox reactions (e.g. Aox4, lyd), immune responses (e.g. *Cd4, Spn*), lipid metabolism (e.g. *Lpcat2, Ffar4*), and cytoskeleton integrity (e.g. *Mld1, Nphs2, Pcsk5, Smpx*). This suggests that glyburide can target and reverse expression of DEGs. To determine whether these glyburide-targetable genes are also differentially expressed in human AD brains, we then explored whether the 23 DEGs were also differentially expressed in the cortex from human AD brains. 6 of these DEGs were confirmed in RNAseq data from control and AD brains (Mayo RNAseq AMP-AD database). These DEGs were involved in signal transduction (e.g. *FFAR4, ECT2L*), proteolysis (e.g. *METAP1D, ADAM18*), gene expression (e.g. *DNMT3A*), and redox reactions (e.g. *IYD).* Together, this suggests that glyburide treatment reverses DEGs associated with signal transduction and excitability due to AD-related pathology.

**Figure 8.**
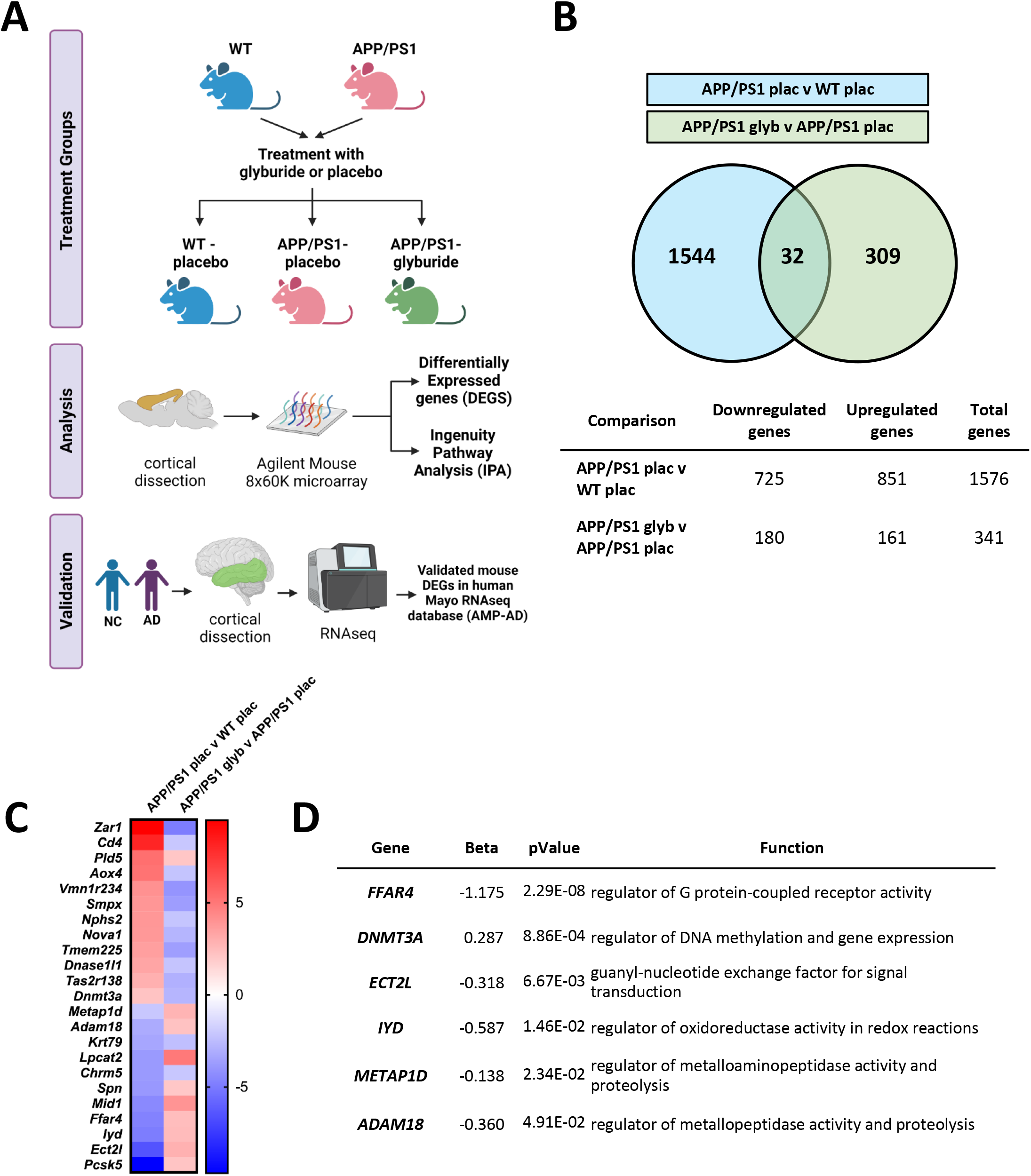
Glyburide treatment reverses disease dependent changes gene expression in the cortex of mice and humans. (A) Venn diagram representing differential gene expression in the cortex of 1) APP/PS1 mice compared to WT controls and 2) glyburide treated APP/PS1 mice compared to placebo treated APP/PS1 control (n=4 mice/group). (B) Heat map highlighting the 32 differentially expressed genes altered due to disease (e.g. APP/PS1 placebo v WT placebo) and subsequently reversed by treatment (e.g. APP/PS1 glyburide v APP/PS1 placebo). (C) Of the genes differentially expressed in mice due to disease and normalized with treatment, a subset of these genes are also altered in the human brain from AD patients. These genes are involved in G-protein coupled receptor activity, signal transduction, redox reactions, neuroinflammation, and proteolysis.

To further clarify how the differential expression of genes related to our functional and pathological findings, we performed Ingenuity Pathway Analysis (IPA). Three main pairwise comparisons were performed to determine the effects of glyburide treatment in APP/PS1 mice: 1) APP/PS1 placebo vs. WT placebo, 2) APP/PS1 glyburide vs. APP/PS1 placebo, and 3) APP/PS1 glyburide vs. WT placebo. These three comparisons identify pathways that were aberrantly affected by disease (APP/PS1 placebo vs. WT placebo), the effects of treatment on disease (APP/PS1 glyburide vs. APP/PS1 placebo), and what pathways are reversed by treatment (meta-analysis of all 3 comparisons). First, 54 pathways were altered by genotype, with 53 showing a positive z-score, or activation, in APP/PS1 mice compared to WT (Figure 9A). Next, 18 pathways displayed a negative z-score in glyburide-treated compared to placebo-treated APP/PS1 mice. Lastly, IPA analysis of APP/PS1 glyburide-treated vs. WT placebo animals shows that 46 pathways remained altered at the transcript level following glyburide treatment in APP/PS1 compared to WT. Interestingly, we identified 3 pathways that were upregulated by disease and downregulated with glyburide treatment to WT levels, suggesting normalization of disease-associated gene expression with glyburide treatment (Figure 9B). These three pathways included: 1) Cardiac Hypertrophy Signaling, 2) Gaq Signaling, and 3) Type II Diabetes Mellitus Signaling (Figure 8C), consistent with the finding that glyburide may reverse vascular remodeling and dysfunction (Figures 6&7). Activation of the Cardiac Hypertrophy Signaling pathway involves G-coupled protein receptor and calcium signaling to alter hemodynamics and vascular remodeling. Consistent with this, the Gaq signaling pathway was also upregulated in the APP/PS1 mice and reversed by glyburide treatment. This pathway is composed of G-coupled protein receptor signaling, including the IP3-DAG-calcium pathway activated by adrenergic receptors, to mediate vasoreactivity and vessel hypertrophy which is known to be mediated by Kir channel activity [39–42]. Lastly, DEGS involved in the Type II Diabetes Mellitus (T2DM) pathway are associated with voltage gated calcium channel activity, protein kinase activity, inflammation, and apoptosis (Figure 9C).

**Figure 9.**
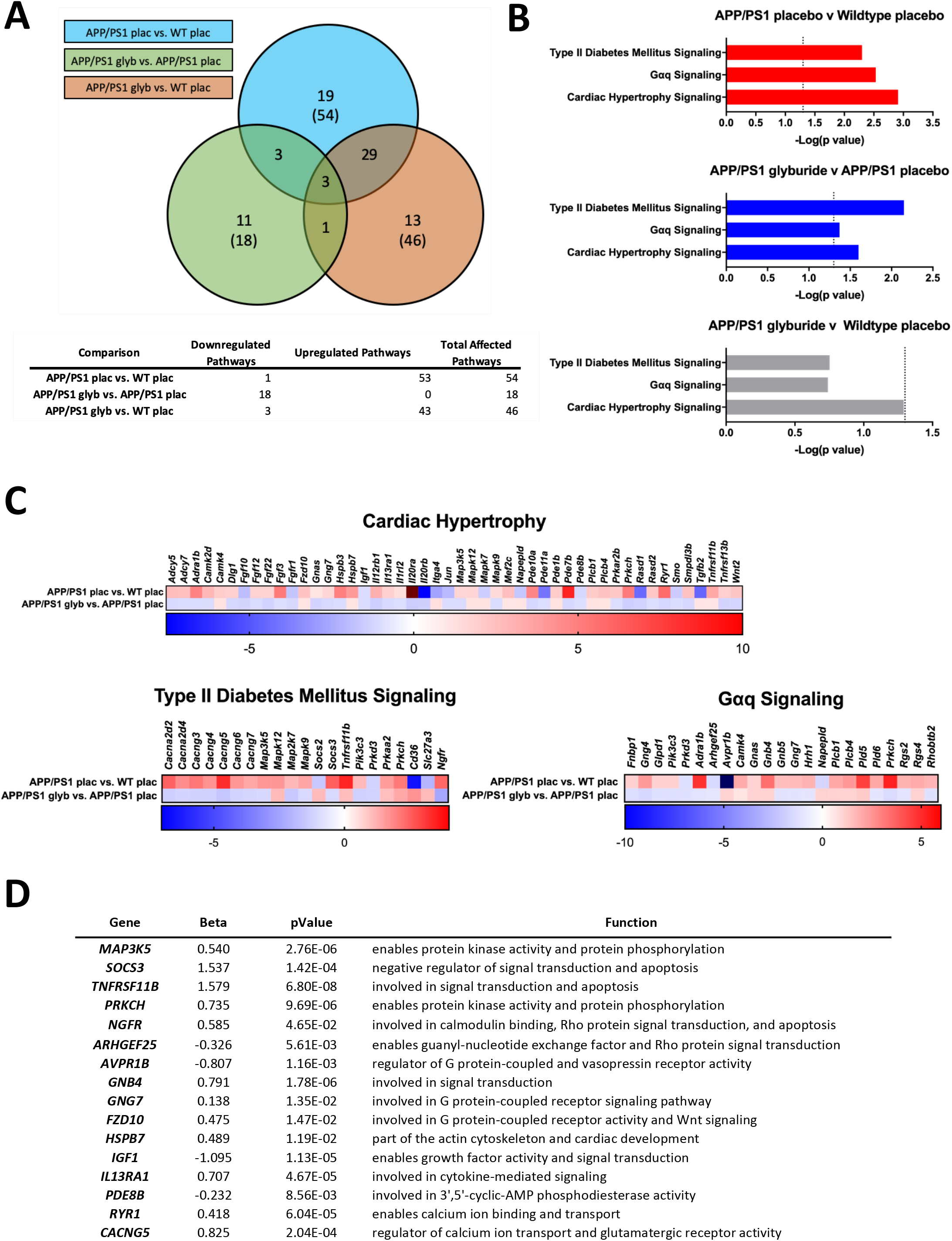
Ingenuity pathway analysis (IPA) identified canonical pathways regulated by APP/PS1 genotype and responsive to glyburide treatment. (A) Venn diagram representing pathways affected by 1) APP/PS1 genotype compared to WT controls, 2) glyburide treatment compared to placebo treatment in APP/PS1 mice, and 3) glyburide treated APP/PS1 mice compared WT controls. APP/PS1 genotype upregulated 53 pathways while glyburide treatment downregulated 18 pathways. Treatment with glyburide restored 3 pathways to wildtype levels. (B) Type II Diabetes Mellitus Signaling, Gαq Signaling, and Cardiac Hypertrophy Signaling Pathways were upregulated due to APP/PS1 genotype (red), downregulated due to glyburide treatment (blue), and indistinguishable from WT (gray). (C) Genes differentially expressed in Type II Diabetes Mellitus, Gαq, and Cardiac Hypertrophy Signaling Pathways upregulated in APP/PS1 mice and subsequently reduced in glyburide treatment compared to placebo. (D) Of the genes identified in Fig 9c in mice, a subset of these genes is also altered in the human brain from AD patients. These genes are largely involved in calcium signaling and G-protein coupled signal transduction suggesting glyburide alters excitability within the brain.

Using the RNAseq data from control and AD brains (Mayo RNAseq AMP-AD database), we explored whether the DEGs associated with these 3 pathways were also differentially expressed in cortex from human AD brains. We identified 16 specific genes that were implicated in human AD and that were normalized by glyburide treatment in APP/PS1 mice. Of these 16 genes, DEGs were enriched in pathways involved in GPCR signaling, calcium signaling, apoptosis, inflammation, and neurotransmission (Figure 9D). Finally, 3 additional pathways were upregulated in APP/PS1 mice compared to WT and downregulated with glyburide treatment, but remained dysregulated compared to wildtype controls: 1) Role of NFAT in Cardiac Hypertrophy, 2) Corticotropin Releasing Hormone Signaling, and 3) Signaling by Rho Family GTPases. These pathways reinforce our findings that systemic glyburide treatment affects vascular K_ATP_ channel activity to impact vascular structure and function, neuronal activity, and AD-related pathology.

### Glyburide treatment reverses arterial stiffness and improves learning and memory

Ingenuity pathway analysis (IPA) implicated cardiac hypertrophy and vascular remodeling as potential targets for glyburide in APP/PS1 mice. Therefore, we explored whether 9-month-APP/PS1 mice with amyloid plaque pathology had any alterations in cardiac function and arterial stiffness, and whether short term treatment with glyburide might reduce arterial stiffness (Figure 9). Echocardiography indicated similar heart rate, cardiac output, E/e’ ratio, ejection fraction, fractional shortening and LV posterior thickness in WT-placebo, APP/PS1-placebo and APP/PS1-glyburide treated mice (Figure 9B), demonstrating that APP/PS1 did not have an overt cardiac phenotype at 9 months of age. Moreover, short term glyburide treatment in the APP/PS1 mice had no adverse effect on cardiac function. In contrast, there was a trend towards higher pulse wave velocity in APP/PS1-placebo mice (8.7 ± 1.2 mm/ms; n = 5) compared to wild type-placebo mice (5.5 ± 1.0 mm/ms; n = 7), suggestive of increased arterial stiffness (Figure 10C). Interestingly, there was a trend towards decreased pulse wave velocity in glyburide-treated APP/PS1 mice (4.9 ± 1.1 mm/ms; n = 5), suggesting a normalization of this phenotype with glyburide treatment (Figure 10C). Higher pulse wave velocity is indicative of arterial stiffness and correlates with changes in cerebrovascular function [43–46]. This suggests the possibility that glyburide reversal of cerebrovascular dysfunction in APP/PS1 mice might underlie the cerebrovascular morphology changes observed with CD31/CD13 staining above.

**Figure 10.**
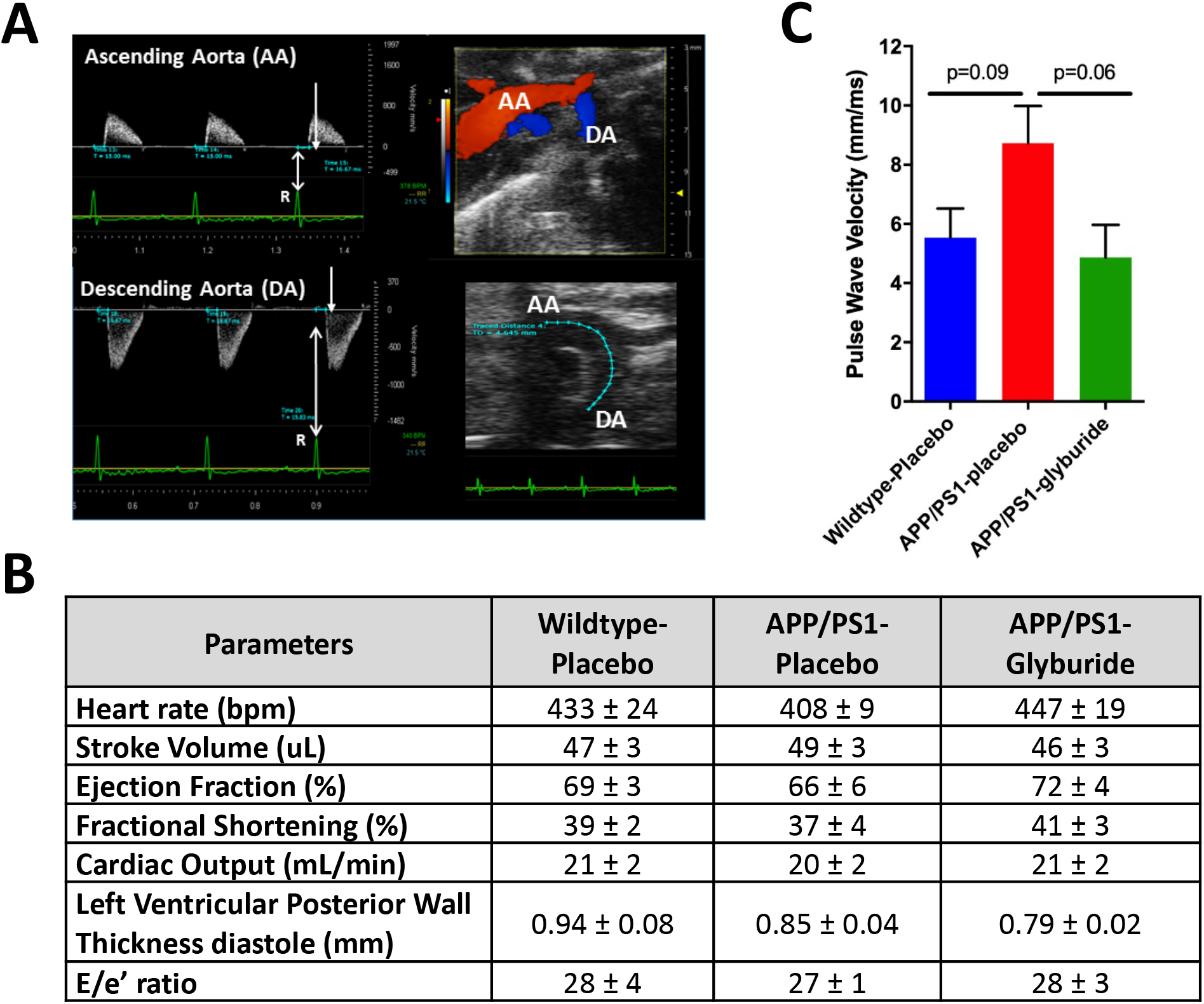
Cardiac function and arterial stiffness in wildtype and APP/PS1 mice following glyburide treatment. (A) Sample image of parasternal short-axis along with 2-D mode images used to assess cardiac function and pulse wave velocity. (B) Cardiac function was comparable between wildtype-placebo, APP/PS1-placebo and APP/PS1-glyburide treated mice on all metrics assessed (n=5-7 mice/group). (C) Pulse wave velocity, a metric of arterial stiffness, was assessed in wildtype-placebo, APP/PS1-placebo and APP/PS1-glyburide treated mice. APP/PS1 mice had a trend towards increased arterial stiffness compared to wildtype. Short term treatment with glyburide decreased pulse wave velocity and reduced arterial stiffness in APP/PS1 mice. Significance determined by one-way ANOVA with Tukey’s test. p values reported to demonstrate trends.

We also investigated whether short-term glyburide treatment in older (8 mo.) APP/PS1 mice might affect learning and memory or brain network functional connectivity (fc) (Fig 11). On the Morris Water Maze behavioral test of learning and memory, glyburide treatment trended towards improvement compared to placebo (Fig 11A-C); both glyburide- and placebo-treated APP/PS1 mice demonstrated learning by decreasing the latency to platform over the 5-day testing period (Fig 11D), and both spent more time in the target quadrant than other quadrants (Fig 11B). Interestingly, glyburide–treated mice showed a trend towards increased platform crossings compared to placebo treated controls (Fig 11C), suggesting improved memory. Functional connectivity intrinsic signal imaging (fcOIS), to assess brain network function connectivity, revealed no differences between wildtype, placebo treated APP/PS1, and glyburide treated APP/PS1 mice at this time point (Fig 11D). This suggests that neither the disease nor therapy negatively affected network connectivity, and together with the above data, suggests that short term glyburide treatment can partially reverse functional deficits in APP/PS1 mice with established amyloid plaque pathology, without adverse effects.

**Figure 11.**
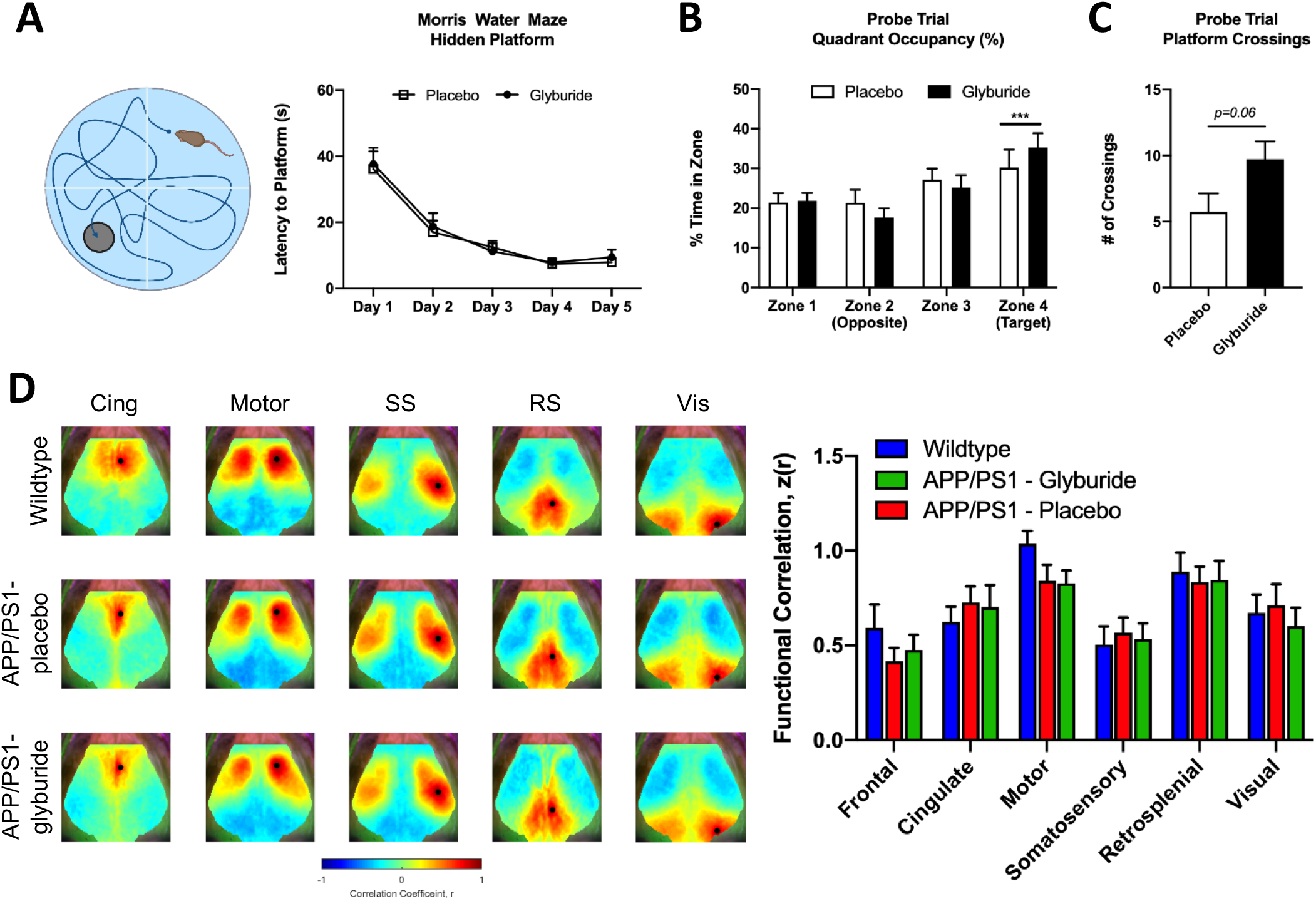
Glyburide treatment improved learning and memory in APP/PS1 mice. (A) Both placebo and glyburide groups demonstrated the ability to learn on the Morris Water Maze (MWM) test by decreasing latency to platform over consecutive testing days (p<0.0001; n=8 mice/group). (B) Both placebo and glyburide groups spent more time in target quadrant during probe trial (p<0.004). (C) Glyburide treated APP/PS1 mice had increased platform crossings (p<0.06) compared to placebo treated controls, suggesting improved learning and memory. (D) No change in functional connectivity was found in the cingulate (Cing), motor (Motor), somatosensory (SS), retrosplenial (RS), and visual (Vis) cortices either due to disease or treatment (n=5-7 mice/group). Significance determined by two-way ANOVA with Tukey’s test. p values reported to demonstrate trends.

## DISCUSSION

### Altered vascular K_ATP_ channel function in AD

K_ATP_ channels are present in neurons and in vascular cells and are important links between cell metabolism and excitability [12], but their role in AD pathology has largely been unexplored. Our initial finding that the vascular-expressed isoforms (*KCNJ8, ABCC9*), but not the neuronal (*KCNJ11, ABCC8*) are specifically upregulated in AD, provides a first indication that vascular K_ATP_ channels may be an important, targetable component of AD. Our studies show that peripheral administration of glyburide decreased amyloid burden and stimulus driven neural and hemodynamic activity, which ultimately reduced Aβ aggregation. Given that glyburide does not efficiently cross the BBB and that vascular K_ATP_ channels regulate vasoreactivity, we hypothesized that enhanced vascular K_ATP_ activity in AD-confirmed by transcriptomic data-might alter the structure and function of the cerebrovasculature. Glyburide-induced reversal of vascular remodeling and transcriptomic changes associated with vessel hypertrophy, as well as arterial stiffness, suggest that K_ATP_ channels play a critical, previously unappreciated, and targetable role linking Aβ with neurovascular dysfunction in AD (Fig 12).

**Figure 12.**
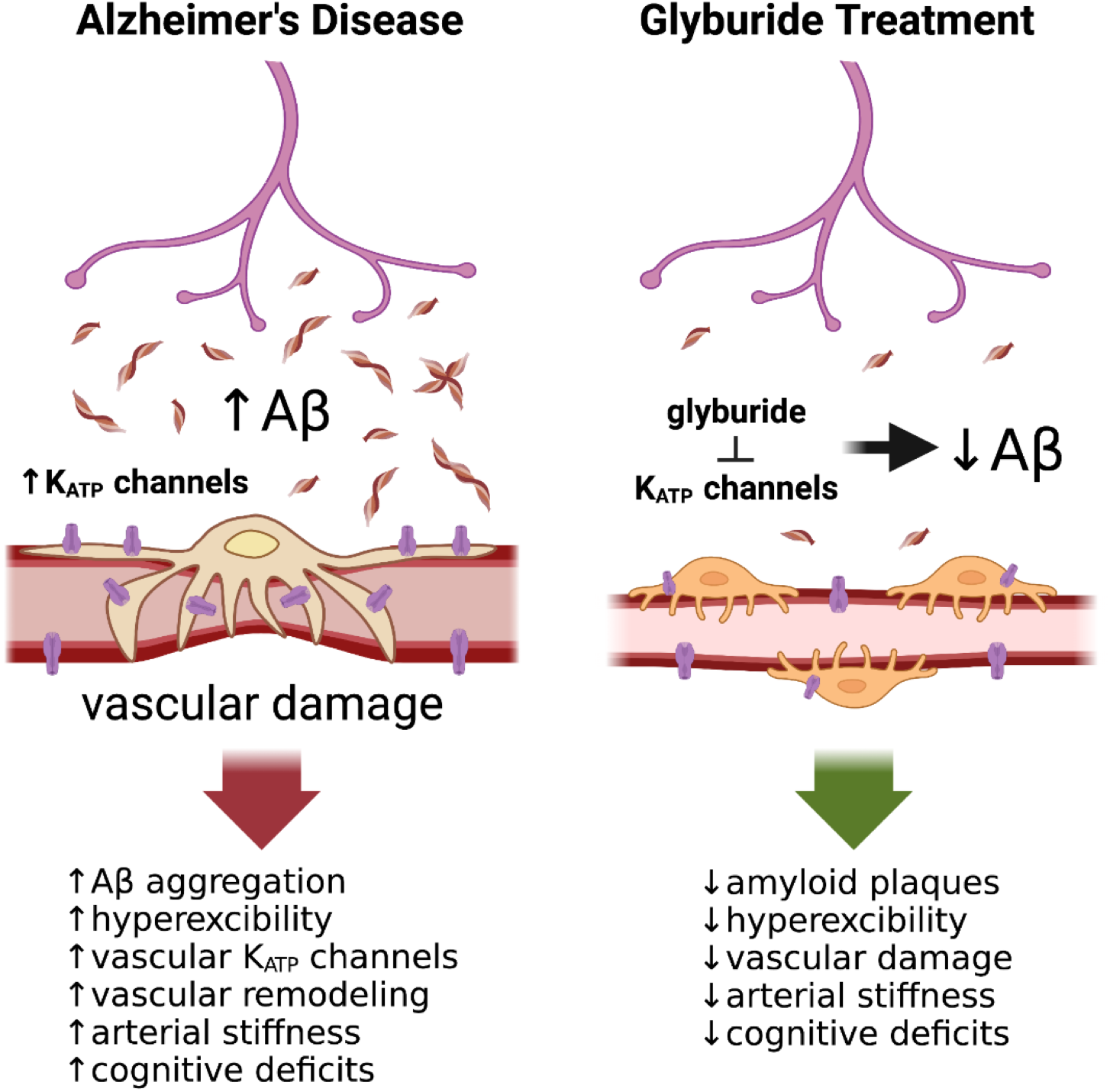
K_ATP_ channel inhibition with systemic glyburide reduces Aβ pathology and vascular damage. Increased Aβ oligomers and amyloid plaques contribute to vascular damage in AD. This coincides with increased expression of vascular K_ATP_ channels in the AD brain, which are important modulators of vasoreactivity. We demonstrate that peripheral treatment with the sulfonylurea and K_ATP_ channel inhibitor, glyburide, reduced the aggregation and activity-dependent production of Aβ. Our data suggest that glyburide targets vascular K_ATP_ channel activity to reduce arterial stiffness, improve vasoreactivity, and normalize pericyte-endothelial cell morphology, providing evidence that vascular K_ATP_ channels are an important, targetable component of Alzheimer’s related pathology.

### Vascular K_ATP_ channels modulate the activity dependent release of Aβ and amyloid plaque formation

In this study, reductions in Aβ deposition and amyloid plaque load were due to changes in Aβ production, not Aβ clearance. One of the most potent drivers of Aβ release is increased neuronal activity [2, 10, 37, 38, 47]. Previous studies demonstrate that the bidirectional modulation of neuronal activity correlates with Aβ levels in the hippocampal ISF. Moreover, if neuronal activity is chronically increased or decreased, via stimulation, deprivation, [10, 37] or modification to sleep [4, 48, 49], amyloid plaque burden is altered in concert. Therefore, the formation of amyloid plaques is both activity- and concentration-dependent. In this study, we explored whether peripheral glyburide treatment decreased neuronal activity and Aβ plaque load. Using electrical and pharmacological stimulation paired with EEG recordings, our work illustrates that the evoked response in the glyburide-treated group was diminished compared to placebo control when presented with the same stimulus. We further demonstrated that reductions in neuronal activity also decreased Aβ production. This effect was surprising because studies in rodents [29], non-human primates [50], or humans [51] show that glyburide does not efficiently cross the blood brain barrier (BBB). In fact, administration of glyburide directly to the CNS prompts neuronal K_ATP_ channel closure and increases neuronal activity, leading to increased ISF Aβ levels [7, 14]. This strongly argues that the paradoxical reduction in Aβ production and aggregation by glyburide is not due to direct inhibition of neuronal K_ATP_ channels, but rather on vascular K_ATP_ channels. Previous work shows that vascular K_ATP_ channels play a major role in the cardiovascular system, where they regulate vasodilation, vasoconstriction, and vascular tone [12, 13, 52, 53]. Opening and closing of K_ATP_ channels triggers vasodilation and vasoconstriction, respectively, which is central to the neurovascular response and necessary to sustain periods of heightened neuronal activity. We demonstrate that the evoked hemodynamic response was dampened in the glyburide-treated APP/PS1 mice compared to placebo, explaining attenuated cerebral blood flow (CBF) and oxy-hemoglobin in the glyburide group when presented with the same stimulus as the placebo group. This suggests that glyburide affects the neurovascular unit in a manner that reduces the activity-dependent release of Aβ. Traditionally, neurovascular coupling (NVC) describes how changes in local neural activity drive changes in local cerebral blood flow [54]. Because neurons do not contain their own energy stores, they rely on the vasculature to provide energy substrates (e.g. oxygen and glucose) to meet local metabolic demand to sustained neuronal activity [55–57]. However, we hypothesize that glyburide acts through the less explored concept of vasculo-neural coupling (VNC) – where changes in vascular tone feed forward to alter neural activity and Aβ release [58–61]. It is well established that K_ATP_ channel activation and inhibition contribute to vascular tone. Our data demonstrate that glyburide treatment reduces arterial stiffness and increases distensibility in a mouse model of Aβ overexpression. Although it is unclear whether glyburide alters vasoreactivity both during a task and at rest, existing evidence supports the idea that changes in vascular K_ATP_ channel activity could restore vascular tone and feedforward to decrease the activity dependent release of Aβ [62–65].

### Implications for future AD treatment

This study is provocative for several reasons. First, it suggests that targeting the vasculature, in this case vascular K_ATP_ channels, holds therapeutic promise for AD. Traditional dogma suggests that CNS therapeutics for AD must cross the BBB in order to be efficacious. In fact, one of the biggest hurdles for treating CNS diseases is effective BBB transport and targeted delivery of therapeutics to vulnerable brain regions. Our study adds to the growing body of evidence that a mechanistic approach to targeting vascular dysfunction holds therapeutic promise in AD.

Secondly, our study supports the evolving treatment strategies could be beneficial in AD despite alterations in CBF. It is well established in humans and rodent models that vascular dysfunction, including decreased CBF, is a pathological hallmark in AD [66–73]. It is also well established that evoked changes in CBF exceed local oxygen demand from increased neuronal activity [74] and changes in the blood oxygen level-dependent (BOLD) signal do not always reliably correlate with neuronal activity [75]. Thus, reducing stimulus-evoked hemodynamic activity does not necessarily equate to reduced neuronal activity or impaired cerebral metabolism. In fact, our data shows the opposite-oxygen consumption, oxygen extraction fraction, and cerebral metabolism are preserved (Fig 6). Since glyburide also reduced arterial stiffness, normalized pericyte-endothelial cell morphology, and improved memory, systemic glyburide treatment was beneficial to functional outcomes. Further studies are needed to fully understand the complex relationship between K_ATP_ channel activity, the neurovascular response, and Alzheimer’s-related pathology.

Second, this study is provocative because it supports the idea that K_ATP_ channels are present in vascular cells in addition to neurons [12, 13] and both of which can be activated or inhibited to be beneficial in AD [76–78]. For example, diazoxide, a K_ATP_ channel agonist, hyperpolarizes multiple neuronal subtypes to reduce neuronal excitability and Aβ/tau release when delivered directly to neurons or the CNS [76]. However, when the K_ATP_ channel antagonists, glibenclamide [77] or tolbutamide [78], are administered orally, depression, anxiety, and memory deficits are reduced following the intracerebroventricular administration of Aβ oligomers [77]. Collectively, these studies demonstrate the following: 1) the route of drug administration dictates whether vascular or neuronal K_ATP_ channels are modulated and 2) inhibiting vascular K_ATP_ channels or activating neuronal K_ATP_ channels both reduce AD-related pathology and behavioral deficits by modulating aberrant cellular excitability. Together, this demonstrates an underappreciated role for K_ATP_ channel activity in the pathogenesis of Alzheimer’s disease.

In our model, we hypothesize that increased Aβ production and aggregation causes vascular remodeling [68, 70], potentially as a consequence of increased vascular K_ATP_ activity, similar to what is seen in Cantu syndrome. Cantu syndrome arises from specific gain-of-function mutations in *KCNJ8* or *ABCC9* [17, 18, 53, 79] and is characterized by chronic vasodilation and decreased vascular tone. This leads to impaired vasoreactivity, and other cardiovascular abnormalities. Importantly, these cardiovascular perturbations can be reversed by glyburide treatment in rodents [17] and potentially in humans [80], consistent with what is observed in our preclinical study with AD-related pathology.

Taken together, we suggest that increased oligomeric Aβ results in K_ATP_ channel activation and vascular dilation, perhaps similar to Cantu syndrome. K_ATP_ channel inhibition with glyburide can reverse these defects, reducing the progression of Aβ related pathology in AD. Our studies demonstrate a critical and previously unappreciated role of K_ATP_ channels in linking neurovascular function and Aβ aggregation in Alzheimer’s disease. Our data further suggest a direct, potential therapeutic approach – targeted inhibition of vascular K_ATP_ channels - to delaying the onset and progression of cognitive impairment in those with underlying Aβ pathology.

## METHODS

### Gene expression alternations in AD brains

To determine whether K_ATP_ channel genes were differentially expressed in AD brain tissue, we analyzed gene expression of pleiotropic genes in publicly available datasets. We analyzed gene expression data from the Mayo Clinic Brain Bank (Mayo) RNAseq study from the Accelerating Medicines Partnership – Alzheimer’s Disease (AMP-AD) portal: the temporal cortex of 80 control, 82 AD, 84 progressive supranuclear palsy (PSP), and 29 Pathologic Aging (PA) brains (syn3163039; syn5550404). Multi-variable linear regression analyses were performed using CQN normalized gene expression measures and including age at death, gender, RNA integrity number, brain tissue source, and flowcell as biological and technical covariates [81].

### APPswe/PS1ΔE9 mice

4-month-old female mice hemizygous transgenic mice for mutations in amyloid precursor protein (APP) and presenilin 1 (PSEN1), the APPswe/PS1ΔE9 mice (1) (APP/PS1) or wildtype littermates on a B6C3 background were used for studies described below. For pulse wave velocity, behavioral, and functional connectivity studies, mice were 9 months of age. Mice were given food and water ad libitum and maintained on a 12:12 light/dark cycle.

### Drug pellet implantation

Glyburide or placebo pellets at a concentration of 2.5mg per 90-day release were purchased from Innovative Research of America (Sarasota, FL) and implanted subcutaneously as previously described [36]. Briefly, 4-month-old APP/PS1 mice were anesthetized with isofluorance, the skin on the lateral side of the neck lifted, and pellets implanted subcutaneously using a stainless steel precision trochar (Innovative Research of America, Sarasota, FL). Based on previous studies, continuous release pellet equates to 30μg/day (2.5mg/pellet; 30μg/day)[29, 36]. For pulse wave velocity, behavioral, and functional connectivity studies, mice were pelleted at 6.5 months and testing performed 2 month later at 8.5 months of age.

### In vivo microdialysis

Guide cannulas (BR-style, BIoanalytical Systems) were stereotaxically implanted into the hippocampus (from bregma, A/P: −3.1mm, M/L: −2.5mm, D/V: −1.2mm at 12° angle) and secured into place with dental cement. Cresyl violet staining was used to validate probe placement post-mortem. Mice were transferred to Raturn sampling cages (Bioanalytical Systems) and microdialysis probes (2 mm; 38 kDa molecular weight cut-off; BR-style, Bioanalytical Systems) inserted into the guide cannula, connected to a syringe pump and infused with artificial cerebrospinal fluid (aCSF; 1.3mM CaCl2, 1.2mM MgSO4, 3mM KCl, 0.4mM KH2PO4, 25mM NaHCO3 and 122mM NaCl; pH=7.35) at a flow rate of 1 μl/minute. Hourly collection of hippocampal ISF began after an 8 hour acclimation period. Steady state levels of ISF glucose, lactate, and Aβ1-x were assayed as described below.

### Drug administration via reverse microdialysis

To determine the clearance rate of ISF Aβ and its half-life in the ISF, Compound E (200μM; Enzo Life Sciences), a gamma secretase inhibitor, was infused directly into the hippocampus via reverse microdialysis as previously described (3). Hourly fractions of ISF were collected and Aβ1-x was measured to calculate half-life as previously described. To determine the activity dependent release of Aβ, picrotoxin (PTX; 25um) was administered via reverse microdialysis, hourly ISF fractions were collected and Aβ1-x measured as described below. Statistical significance was determined using a one-way ANOVA and Dunnett’s multiple comparison post-hoc test. Data is represented by means +/− SEM.

### Tissue processing for post-mortem analyses

7-month-old APP/PS1 mice were anesthetized with an overdose of sodium pentobarbital and transcardially perfused with heparinzed ice-cold phosphate buffered saline (PBS). The brains were removed and dissected out. One hemisphere fixed in 4% paraformaldehyde for 48 hrs at 4°C, and cryoprotected in 30% sucrose for immunohistochemical procedures. Brains were sectioned on a freezing microtome at 50μm. Serial sections (300μm apart) through the anterior-posterior aspect of the hippocampus were used for immunostaining and quantification. Brain regions from the contralateral hemisphere, including the anterior cortex, posterior cortex, hippocampus, striatum, cerebellum, hypothalamus, and septum, were dissected out and flash frozen. Specific brain regions were further processed for ELISAs, western blot, and qPCR. For ELISAs, hippocampal tissue was hand homogenized in 10x w/v RIPA buffer with protease and phosphatase inhibitors (Roche). Homogenates were centrifuged at 14,000 rpm for 30 minutes at 4°C. Supernatant was collected and used as tissue lysate. Total protein concentration in lysates was determined by Micro BCA (ThermoScientific) according to manufacturer’s instructions.

### Immunohistochemical and quantification procedures for Aβ, X34, GFAP, CD68, Iba1, CD31, and CD13 staining

Serial sections (300μm apart) through the anterior-posterior aspect of the hippocampus were immunostained for either Aβ deposition or X-34. For Aβ deposition, free floating sections were stained using a biotinylated, HJ3.4 antibody (anti-Aβ1–13, mouse monoclonal antibody generated in-house) and developed using a Vectastain ABC kit and DAB reaction. For fibrillary plaques, free floating sections were permeabilized with 0.25% Triton X-100 and stained with 10 μM X-34 in 40% ethanol + 0.02M NaOH in PBS [82]. For GFAP, CD68, Iba1, CD13, and CD31x staining, free floating sections were blocked with 3% donkey serum and incubated overnight with either rabbit anti-GFAP (1:10000, Dako), rat anti-CD68 (1:200, BioRad), or rabbit anti-Iba1 (1:5000, Wako). Stained brain sections were imaged using a NanoZoomer slide scanner (Hamamatsu Photonics) and the percent area occupied by HJ3.4 or X-34 was quantified by a blinded researcher as previously described[7, 83]. Statistical significance was determined using a two-tailed, unpaired t-test. Data is represented by means +/− SEM.

### Aβ_1-X_, Aβ_40_, and Aβ_42_ ELISA

For total Aβ concentrations within the hippocampal ISF, Aβ1-x was quantified using a monoclonal capture antibody targeted against Aβ13-28 (m266) and a biotinylated detection antibody targeted against Aβ1-5 (3D6), both generous gifts from Dr. Ron DeMattos, Eli Lilly and Co., Indianapolis, IN. After incubation with streptavidin-poly-HRP-20, the assay was developed using Super Slow TMB (Sigma) and the plates read on a Bio-Tek Synergy 2 plate reader at 650nm. For Aβ40 and Aβ42 levels, the soluble and insoluble fractions of hippocampal tissue homogenates were isolated and assayed using sandwich ELISAs as previously described[7, 82]. Briefly, Aβ40 & Aβ42 were quantified using monoclonal capture antibodies targeted against amino acids 45-50 (HJ2) or 37-42 (HJ7.4), respectively. For detection, both Aβ40 and Aβ42 used a biotinylated monoclonal antibody against the central domain (HJ5.1B), followed by incubation with streptavidin-poly-HRP-40. Assays were developed using Super Slow TMB (Sigma) and the plates read on a Bio-Tek Synergy 2 plate reader at 650nm.

### Glucose and lactate measurements

Glucose and lactate measurements from blood and hippocampal ISF were quantified using an YSI 2900 analyzer as previously described [7, 84].

### Insulin ELISAs

Serum was collected after 5 hours of fasting and insulin was measured by ELISA (Alpco: Cat. # 80-INSMSU-E01; 80-CPTMS-E01) according to manufacturer’s instructions[83].

### Insulin degrading enzyme (IDE) activity assay

IDE activity was measure in cortical homogenates posttreatment for1 hour using the SensoLyte IDE Activity Assay Kit (AnaSpec, Inc: Cat. # AS-72231) according to manufacturer’s instructions. Frozen cortex was hand homogenized 5x w/v in assay buffer, centrifuged, and the supernatant used as lysate. Fluorescence was measured every 10 minutes for 90 minutes and the relative fluorescence at 90 minutes per mg of protein (determined by MicroBCA) was quantified.

### qPCR analysis

RNA isolation from mouse brain was performed as previously described [85]. Briefly, anterior cortex was homogenized by trituration through a 23-gauge needle in TRIzol (Invitrogen). Chloroform (1:5) was added, samples mixed, then centrifuged at 13,000 g for 15 minutes at 4°C. Chloroform was removed, diluted 1:1 in 70% ethanol, and purified using RNeasy columns and reagents (QIAGEN). RNA concentration was measured using a NanoDrop spectrophotometer, and reverse transcription was performed using a high-capacity RNA-cDNA kit (Applied Biosystems [ABI]) with 1 μg RNA per 20 μl reaction. Real-time qPCR was performed with ABI TaqMan primers and reagents on an ABI Prizm 7500 thermocycler according to the manufacturer’s instructions. All mRNA measurements were normalized to β-actin mRNA levels.

### Western blot analyses for APP metabolism

Cortical tissue was homogenized in 10-fold volume lysis buffer (10 mM Tris-HCl, pH 8.0, 1 mM EDTA, 1% Triton X-100, 0.1% SDS, and 150 mM NaCl) and centrifuged at 15,000 g at 4°C. 50 μg of protein were loaded following quantification with a BCA assay onto a 4–20% Bis-Tris gel and transferred to nitrocellulose membrane. Full-length APP was detected using 6E10 (1:1,000; Biolegend), and C99 fragment was detected using 82E1 (1:1,000; IBL America). Protein detection was visualized using an HRP-coupled anti–mouse secondary antibody (1:5,000) and ECL reagent. Images of the blot were captured using a G Box (SynGene) and analyzed in ImageJ[82].

### Forepaw stimulation

Needle electrodes were inserted into the dorsal and ventral sides of the left and right forepaws between digits 2 and 3. The stimulation paradigm consisted of 5 seconds of rest, followed by a 10 second stimulus train, then 45 seconds of rest administered in a block design (15 trials per mouse). Electrical stimuli were 0.3ms pulses delivered at 3Hz at an amplitude of 1mA driven by a constant current stimulus isolation unit (World Precision Instruments).

### Functional neuroimaging

Six weeks post glyburide or placebo pellet implantation, mice were imaged using interleaved optical intrinsic signal imaging (OISI) and laser speckle contrast imaging (LSCI) under Ketamine/Xylazine anesthesia (Ketamine: 86.9mg/kg; Xylazine: 13.4mg/kg) as previously described [86–88]. Temporal multiplexing of these modalities allows for simultaneous imaging of relative changes in oxygenated (HbO) deoxygenated (HbR), and total (HbT) hemoglobin, and cerebral blood flow (CBF). All mice were imaged in 2 separate sessions separated by 1 week: a stimulus session in which mice were imaged for 15 minutes during forepaw stimulation, and a resting state session during which mice were imaged for 45 min). All imaging data were acquired at 30 Hz. Data from all mice were subject to an initial quality check prior to analysis. Data runs in which reflected light level intensity (mean value over the brain) varied as a function of time by greater than 1% for any wavelength were excluded from further analysis. Evoked hemodynamics were bandpass filtered from 0.009-0.5 Hz, and resting state activity was bandpass filtered from 0.009-0.08 Hz. After filtering, each pixel’s time series was downsampled from 30 Hz to 1 Hz, and all further analysis was performed only on those pixels labeled as brain using a manually-constructed brain mask. Common variance across all brain pixels was removed using global signal regression [89]. Image sequences were coregistered to a common atlas space as previously described[90]. Changes in relative CMRO2 were calculated from the image sequences of CBF, HbO and HbR, and assuming values for the vascular weighting constants and baseline concentrations of HbO and HbR following established protocols [87]. Oxygen extraction fraction (OEF) was estimated as OEF=CMRO2/CBF (i.e. assuming 100% arterial oxygen saturation and 0% venous oxygen saturation). Time courses for each contrast were created by averaging all pixel time courses within 50% of peak response within forepaw somatosensory cortex.

### Resting state functional connectivity

RSFC was performed as previously described [90]. Briefly, fourteen seed locations were positioned at coordinates corresponding to left and right frontal, cingulate, motor, somatosensory, retrosplenial, and visual cortices. Seed time traces were calculated by averaging time traces within 0.25 mm (30 pixels) of a seed locus and correlated with every other pixel in the brain to create a functional connectivity map for that seed. Individual functional connectivity maps were averaged within each cohort. Bilateral connectivity was determined by correlating each seed with its contralateral homolog, thus producing seven bilateral correlation values for each mouse. Pearson r values were converted to Fisher Z measures prior to any averaging or statistical comparison.

### EEG recordings

Six weeks post glyburide or placebo pellet implantation, APP/PS1 mice were subjected to electroencephalography (EEG) recordings in S1FL region of cortex contralateral to forepaw stimulation as previously described[91]. Whiskers are stimulated for 10s on and 50 second off. EEG activity is assessed using a P511K A.C. preamplifier (Grass Technologies), digitized with a DigiData 1440A data acquisition system (Molecular Devices) and recorded digitally using pClamp 10.2 (Molecular Devices). Changes in EEG amplitude and power analysis during the whisker stimulation will be compared to baseline in each animal. Group averages in EEG amplitude during stimulation compared to baseline will be calculated for each group.

### Plexiglass window implantation

Prior to imaging, Plexiglas windows were affixed to mouse skulls as previously described[92]. Briefly, mice were anesthetized with isoflurane (5% induction, 2% maintenance) with their heads secured in a stereotaxic frame, and placed on a heating pad maintained at 37°C (via feedback from a rectal probe). After shaving and cleaning the scalp, a midline incision was made along the scalp, keeping the skull intact. A clear Plexiglas window with pre-tapped holes was fixed with translucent dental cement (C&B-Metabond, Parkell Inc., Edgewood, NY), and a Plexiglas coverslip was screwed onto the imaging window to protect the window from scratching while the mice were in their home cages. Post-surgery, mice were returned to their home cages and allowed to recover for at least 72 hrs. The total mass of the window, screws, and cover was approximately1 g.

### Microarray analysis

Microarray was performed on the anterior cortex from 7 month-wildtype-placebo, APP/PS1-glyburide treated, and APP/PS1-placebo (n=4 mice/group) following transcardial perfusion as previously described [93]. RNA was isolated from flash frozen cortex samples as above and submitted to the Genome Technology Access Center at Washington University for quality control, MessageAmp RNA library preparation with Kreatech ULS, and Agilent Mouse 8×60K microarray. Differentially expressed genes (DEGs) were identified using Bayesian statistics, where significant genes were filtered out with a p value<0.05 or FDR-q<0.05, and a fold-change>/=2. For Ingenuity Pathway

Analysis (QIAGEN, Venlo, Netherlands), fold changes and p-values were imported for all expression values. Networks and pathways were analyzed for significant enrichment of module genes and were ranked by −log p-values. To quantify a significant association between differentially expressed genes and molecular in annotated pathways, a right-tailed Fisher’s exact test was employed [94]. P-values of < 0.05, FDR<0.003 was considered significant.

### Pulse wave velocity and cardiac function

Cardiac function and arterial stiffness were assessed in 9-month-old female wildtype-Placebo, APP/PS1-Placebo and APP/PS1-Glyburide. The thoracic area was treated with hair removal cream before imaging. Transthoracic echocardiography was performed using a Vevo 2100 LAZR ultrasound system (FUJIFILM/VisualSonics, Inc.; Toronto, Canada) equipped with a 30 MHz linear array transducer[95]. Mice were lightly anesthetized (1-1.5% inhaled isoflurane) and animals were placed in temperature controlled platform with two-lead electrodes to monitor heart rate during imaging. Parasternal short-axis along with 2-D mode images were obtained to assess stroke volume, heart rate, cardiac output, posterior wall thickness, ejection fraction, and fractional shortening [96, 97]. Mitral valve early filling velocities (E) and septal annular velocities (e’) were obtained using pulsed Doppler and tissue Doppler, respectively. E/e’ was calculated as an index of left ventricular (LV) filling pressure [96, 98]. To determine arterial stiffness, the pulse wave velocity was calculated as follows: PWV = Distance (D)/Time (T), where D is the distance in mm between the ascending aorta site to descending aorta site and T = (R point of the EKG to the foot of the ascending aorta flow) - (R point of the EKG to the foot of the descending aorta flow) in msec [98]. All images were acquired and analyzed with Fujifilm Visualsonic VevoLab software (Version 3.2.6).

### Morris water maze

Morris water maze (MWM) was performed as previously described [99]. Briefly, the test consisted of four trials (60-second maximum, 15 minute interval) per day for five consecutive days. Escape latency was measured as an average of four trials each day. A probe trial was conducted two hours after the final training session of the fifth day. The visible platform task consisted of four trials each day for two days with the escape platform marked by a visible cue and moved randomly among 4 locations. Trajectories, time spent in each maze quadrant, velocity, and distance moved were recorded using ANY-Maze tracking software (Stoelting Co., Wood Dale, IL).

### Statistics

Statistical analysis using t-tests, one-way ANOVAs, and 2way ANOVAs and the appropriate post-hoc tests were performed for each experiment above. P values equal to or less than 0.05 were considered significant.

### Study Approval

All procedures were carried out in accordance with an approved IACUC protocol from Wake Forest School of Medicine or Washington University School of Medicine.

## AUTHOR CONTRIBUTIONS

SLM and DMH conceived of the study. SLM, DMH, MSR, CGN, AQB, JPC, NCD, CMK, LAC, and DID contributed to study design. SLM, MSS, EEC, WRM, ARB, NCD, CMC, SMD, JG, TEM, JAS, TEO, AQB, LAC, DID, and CMK performed experiments, data analysis, and data interpretation. SLM, CGN, and DMH wrote the manuscript. All authors discussed the results and commented on the manuscript.

## ACKNOWLEDGEMENTS

We would like to acknowledge the following grants: 1K01AG050719 (SLM), R01AG068330 (SLM), BrightFocus Foundation (A20201775S; SLM), Charleston Conference on Alzheimer’s disease New Vision Award (SLM), R01NS102870 (AQB), the McDonnell Center for Systems Neuroscience (SLM and AQB), P01NS080675 (DH, AQB, SLM, JC), R01 HL140024 (CGN), F31AG066302 (CC), T32AG033534 (JG), and T32AA007565 (SD). We would also like to acknowledge the Mayo Clinic Brain Bank (Mayo) RNAseq study from the Accelerating Medicines Partnership - Alzheimer’s Disease (AMP-AD) portal. Study data were provided by the following sources: The Mayo Clinic Alzheimers Disease Genetic Studies, led by Dr. Nilufer Taner and Dr. Steven G. Younkin, Mayo Clinic, Jacksonville, FL using samples from the Mayo Clinic Study of Aging, the Mayo Clinic Alzheimers Disease Research Center, and the Mayo Clinic Brain Bank. Data collection was supported through funding by NIA grants P50 AG016574, R01 AG032990, U01 AG046139, R01 AG018023, U01 AG006576, U01 AG006786, R01 AG025711, R01 AG017216, R01 AG003949, NINDS grant R01 NS080820, CurePSP Foundation, and support from Mayo Foundation. Study data includes samples collected through the Sun Health Research Institute Brain and Body Donation Program of Sun City, Arizona. The Brain and Body Donation Program is supported by the National Institute of Neurological Disorders and Stroke (U24 NS072026 National Brain and Tissue Resource for Parkinsons Disease and Related Disorders), the National Institute on Aging (P30 AG19610 Arizona Alzheimers Disease Core Center), the Arizona Department of Health Services (contract 211002, Arizona Alzheimers Research Center), the Arizona Biomedical Research Commission (contracts 4001, 0011, 05-901 and 1001 to the Arizona Parkinson’s Disease Consortium) and the Michael J. Fox Foundation for Parkinsons Research.

The authors have declared that no conflict of interest exists.

